# Lipid flipping in the omega-3 fatty-acid transporter

**DOI:** 10.1101/2022.05.31.494244

**Authors:** Chi Nguyen, Hsiang-Ting Lei, Louis Tung Faat Lai, Marc J. Gallenito, Doreen Matthies, Tamir Gonen

**Affiliations:** Howard Hughes Medical Institute, University of California Los Angeles, Los Angeles CA 90095; Janelia Research Campus, Howard Hughes Medical Institute, 19700 Helix Drive, Ashburn, VA 20147; Unit on Structural Biology; Division of Basic and Translational Biophysics, Eunice Kennedy Shriver National Institute of Child Health and Human Development, National Institutes of Health, Bethesda, MD 20892; Departments of Biological Chemistry and Physiology, University of California Los Angeles, Los Angeles CA 90095

**Keywords:** Omega-3 fatty-acid, lipid, transporter, membrane protein, cryo-EM

## Abstract

Mfsd2a is the primary transporter for the docosahexaenoic acid (DHA), an omega-3 fatty acid, across the blood brain barrier (BBB). Defects in Mfsd2a are linked to ailments from behavioral, learning, and motor dysfunctions to severe microcephaly. Mfsd2a typically transports long-chain unsaturated fatty-acids, including DHA and α-Linolenic acid (ALA), that are attached to the zwitterionic lysophosphatidylcholine (LPC) headgroup. Even with two recently determined structures of Mfsd2a the molecular details of how this transporter performs the energetically unfavorable task of translocating and flipping lysolipids across the lipid bilayer remained unclear. Here, we report five single-particle cryo-EM structures of the *Danio rerio* Mfsd2a (drMfsd2a): in the inward-open conformation in the ligand-free state and bound to ALA-LPC at four unique positions along the substrate translocation pathway. These Mfsd2a snapshots detail the Na^+^-dependent flipping mechanism of the lipid-LPC from outer to inner membrane leaflet during ligand translocation through the Mfsd2a substrate tunnel and release for membrane integration on the cytoplasmic side. These results also map Mfsd2a mutants that disrupt lipid-LPC transport and are associated with known disease. Together these results provide a model for omega-3 fatty-acid transport and has the potential for the design of the delivery strategies for amphipathic drugs across the BBB.

## Introduction and background

Docosahexaenoic acid (DHA, 22:6) is an important component of cellular membranes where it forms structural and functional interactions with cholesterol and integral membrane proteins^1^ (Fig. 1a). DHA is enriched in the central nervous system (CNS), comprising of 5% of the brain by mass and 20% of all lipids in neuronal cell membranes^2^. Remarkably, the concentration of DHA approaches almost 50% of all lipids in the phospholipid bilayer of neuronal synapses^22^. It is not surprising then that inadequate DHA results in several CNS-related ailments including learning and memory deficits, dyspraxia, and dyslexia^3,4^. Despite its essential roles in CNS structure and function, the brain lacks robust mechanisms to biosynthesize DHA, which is estimated to have high turnover rates on the order of minutes^5^. Instead, the brain acquires DHA from plasma by transport across the blood-brain barrier (BBB)^6^. Although many studies have gleaned important insights into the roles of tight junctions at the BBB^7–9^, recent studies have revealed that DHA is delivered to the brain via selective transport through endothelial cells^5,6^. Unlike transport between cells at the tight junctions, selective DHA transport across the BBB involves capturing the lipid on the plasma side, integrating into metabolic processes and/or transporting through, and finally releasing the molecule to the opposite side of the cell to the brain^5,6^.

**Fig. 1.**
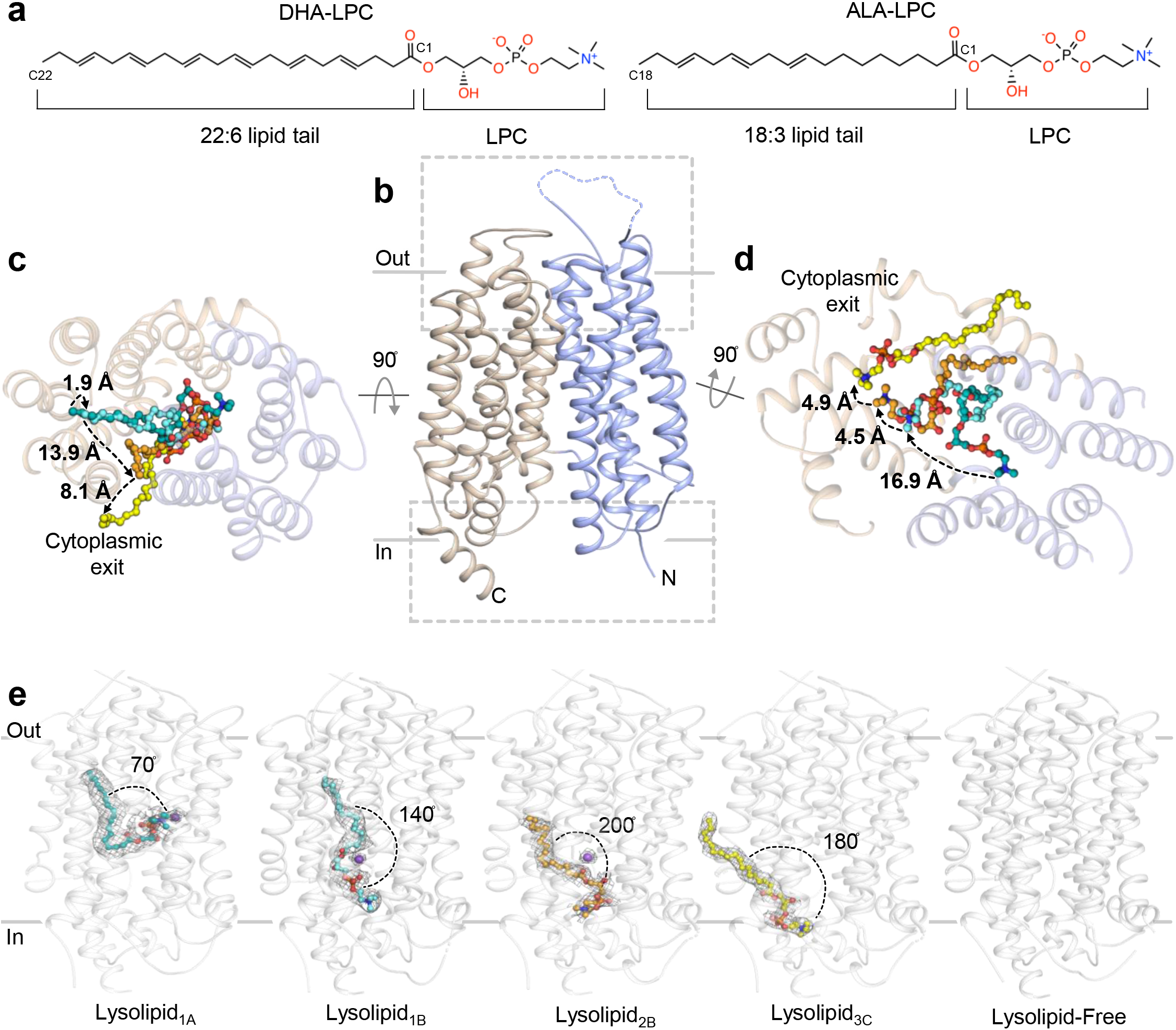
Overall topology and lysolipid transport by drMfsd2a. **a**, The preferred Mfsd2a DHA-LPC substrate and the ALA-LPC ligand observed in the drMfsd2a structures. **b**, Overall architecture of drMfsd2a with the N-terminal domain, TM1-6, in blue and the C-terminal domain, TM7-12, in wheat. **c**, Extracellular view of the substrate translocation pathway with the overlay of the four lysolipid positions observed in drMfsd2a. Dashed lines represent the proposed transport path of the C18 of the lipid tail through a cleft between the N- and C-domains. **d**, Cytoplasmic view of the substrate translocation pathway with the overlay of the four lysolipid positions observed in drMfsd2a through a cleft between the N- and C-domains. Dashed lines represent the proposed transport path of the choline of the LPC headgroup. **b-d**, Ligands illustrated as stick and sphere. **e**, The ligand-free and the four ALA-LPC conformations observed for drMfsd2a starting from the position closest the extracellular towards cytoplasmic side. Protein rendered as grey cartoon. Na^+^ represented as purple spheres. Electron density shown as grey mesh.

The major facilitator superfamily (MFS) is a large class of membrane proteins that span all kingdoms of life having diverse functions and substrate specificities^10,11^. MFS comprise over 25% of all transmembrane proteins and play key roles in physiology^9^. Two of the most well studied MFSs are LacY^12^ and GLUTs (SLC2s)^13^ for importing sugar into cells and for maintaining glucose homeostasis. Although members of MFS share low sequence identity and bind a variety of substrates, they all share a similar twelve transmembrane helix (TM) topology and use the “rocker-switch” mechanism to transport cargos across membranes^11,14^. During the rocker-switch motion, the substrate is initially bound, followed by sequential structural changes that reverberate outward from the ligand binding site, resulting in the switch motion that closes the transporter on one side and opens the transporter on the opposite side of the membrane^11,14^.

The MFS domain containing protein-2a (Mfsd2a) was initially discovered as an orphan receptor in 2008^15^ and later revealed as the primary DHA transporter across the BBB^6^. Importantly, Mfsd2a has been shown to not only transport DHA but is also required for the formation, development, and function of both the BBB and CNS^16–18^. Defects in Mfsd2a disrupt normal brain development and function resulting in phenotypes including anxiety, learning deficits, ataxia, and severe microcephaly^6,16,18–20^., similar to symptoms in animals lacking omega-3 fatty acids. Consistent with these findings, it was recently demonstrated that Zika virus infection causes microcephaly by degradation of Mfsd2a and supplementation of DHA recues this phenotype^21^. In contrast to other members of MFS that shuttle water-soluble ligands such as small-molecule drugs, oxyanions, metabolites, sugars, and amino acids, the Mfsd2a transports fatty acids^6^. Mechanistic studies revealed that Mfsd2a is a symporter for Na^+^ and is selective towards long chain, unsaturated fatty-acids including DHA (22:6), α-Linolenic acids (ALA, 18:3), and oleic acids (18:1) that are covalently attached to the zwitterionic lysophosphatidylcholine (LPC) headgroup, which is comprised of a phosphate and choline moieties^6,22^ (Fig. 1a).

Lipids can move laterally in membranes but because of their bulky, charged headgroups, they cannot easily flip vertically between leaflets of the bilayer. Lipid flipping between the membrane bilayers is thought to be energetically costly because this process not only entails the translocation of the charged headgroup across the hydrophobic acyl-chain region of the bilayer but also the flipping of lipid molecules to align with the opposite membrane leaflet while an electric and hydrophobic seal is maintained. Cells accomplish membrane leaflet asymmetry by specialized transporters termed floppases^23^ and flippases^24–26^. These proteins perform these challenging tasks by coupling lipid transport and flipping with energetically favorable processes such as ATP hydrolysis or shuttling ions down their electrochemical gradients^24–27^. In 2017, studies linked the transbilayer transport of DHA-LPC to Mfsd2a by demonstrating its flippase activity^28^, a process defined as the translocation of lipids from the outer to inner membrane leaflet^26^. The key finding from these studies illustrates pronounced concentration of DHA-LPC in the inner versus the outer leaflet of the membrane in the presence of Mfsd2a. Since most flippases so far are ATP-dependent ABC cassettes, Mfsd2a is one of few characterized flippases outside this super family and the only one known to perform lipid flipping in an ATP-independent and Na^+^-dependent manner^24,28,29^. The three other known MFS lipid transporters are Mfsd2b^30^, LplT^31^, and Spns2^32^; however, they confer different substrate specificities and their transport mechanisms are yet to be defined. To date, even with two recently published structures from mouse in the outward-facing^33^ and chicken in the inward-facing^22^ conformations, the molecular details of how Mfsd2a transports and flips DHA-LPC across the lipid bilayer is obscure. Specifically, knowledge of how Mfsd2a binds, transports, and flips both hydrophobic and charged moieties of lipid-LPC is yet to be fully detailed. Moreover, how Na^+^ contributes to these processes is unclear. As such, the novel ATP-independent and Na^+^-dependent DHA-LPC transport and flippase mechanism of Mfsd2a is yet to be described.

Here, we present five structures of the Zebrafish (*Danio rerio*) isoform B Mfsd2a (drMfsd2a) in the inward-facing conformation determined by single-particle cryo-electron microscopy (cryo-EM) in the ligand-free state and harboring the ALA-LPC lysolipid, a known Mfsd2a substrate^22^, in four unique binding positions (Fig. 1, S1-3, Table S1-2). These four ligand binding configurations allowed the (1) mapping of the transport and flipping of the lysolipid as it traverses through an internal tunnel and (2) the release of the substrate to the cytoplasmic side of drMfsd2a for integration into the inner membrane leaflet (Fig. 1c-e). The details gleaned from these studies provide the molecular models of the proposed drMfsd2a lipid, phosphate, choline, and Na^+^ binding sites, and how these components are orchestrated during the transport and flipping of the lysolipid. Instead of a linear substrate tunnel, our studies illustrate the presence of three connected but distinct compartments that each contain a hydrophobic and a charged cavity to accommodate the lipid tail and LPC of the lysolipid. We demonstrate that the binding of the lysolipid and ligation of Na^+^ within and across these compartments play key roles in lysolipid flipping. We also identified the interactions between the lysolipid and drMfsd2a and map key residues that when mutated (1) disrupt lysolipid transport and (2) are associated with known Mfsd2a -related diseases. Together this study adds to the repertoire of transport mechanisms by MFS by generating models for how members of this superfamily of proteins transport lipid cargos. Given its roles in brain function and development, our studies lay the foundation for better understanding of neurological conditions and motor dysfunctions linked to defects in this family of transporters. Given its ligand preference, there is also potential for exploiting the Mfsd2a mechanism for the design of delivery strategies for hydrophobic and amphipathic drugs across the BBB.

### Overall topology of drMfsd2a in complex with lipids

drMfsd2a is a 59 kDa protein with high sequence and structure homology to Mfsd2a from other species. Small membrane proteins of this size are exceedingly challenging by single-particle cryo-EM^34^. As such, we generated and formed a complex between FAB^35^ and drMfsd2a to facilitate cryo-EM structure determination (Fig. S1-3, Table 1-2). Alignment between the human and drMfsd2a sequences revealed a 76% similarity and 61% identity (Fig. S4). Given that most MFSs share low sequence identities but a conserved fold^10^, sequence conservation between these two homologs suggests that the zebrafish protein can serve as a model for understanding the human transporter.

**Table 1.** Transport activities of Mfsd2a mutants.

| Residue | Mutation | Transport Activity | drMfsd2a Site | DrMfsd2a Lysolipid | drMfsd2a Function |
| --- | --- | --- | --- | --- | --- |
| Tyr50 | Ala<sup>33</sup> | Reduced | Z<sub>A</sub> | Lysolipid<sub>1A</sub> | H-bond |
|  |  |  | Na<sub>B</sub> | Lysolipid<sub>1A</sub> | Na<sup>+</sup> coordination |
| Gln51 | Leu<sup>22</sup> | Reduced | Z<sub>A-B</sub> | Lysolipid<sub>1A, 1B</sub> | H-bond |
|  |  |  | Na<sub>B</sub> | Lysolipid<sub>1A, 1B</sub> | Na<sup>+</sup> coordination |
| Phe59 | Ala<sup>33</sup> | Reduced | Chamber<sub>1</sub> | Lysolipid<sub>1A, 1B</sub> | Lipid tail interaction |
| Arg84 | Ala/Lys/Glu<sup>22</sup> | Abolished | Z<sub>A</sub> | Lysolipid<sub>1A, 1B</sub> | Salt-bridge |
| Asp87 | Ala<sup>33</sup> | Abolished | Na<sub>A</sub> | Lysolipid<sub>1A</sub> | Na<sup>+</sup> coordination |
|  |  |  | Z<sub>A</sub> | Lysolipid<sub>1A</sub> | Salt-bridge |
| His156 | Ala<sup>33</sup> | Abolished | Z<sub>B</sub> | Lysolipid<sub>2B</sub> | H-bond |
| Ser160 | Leu<sup>19, 33</sup> | Reduced | Z<sub>B</sub> | Lysolipid<sub>2B</sub> | H-bond |
| Arg180 | Ala<sup>33</sup> | Abolished | Z<sub>B</sub> | Lysolipid<sub>1B, 2B</sub> | Salt-bridge |
| Met181 | Ala/Phe<sup>22</sup> | Reduced | Chamber<sub>2</sub> | Lysolipid<sub>2B</sub> | Lipid tail interaction |
| Thr182 | Phe/Met<sup>33</sup> | Reduced | Chamber<sub>3</sub> | Lysolipid<sub>3C</sub> | Lipid tail interaction |
| Glu184 | Ala<sup>33</sup> | Reduced | Na<sub>B</sub> | Lysolipid<sub>1B, 2B</sub> | Na<sup>+</sup> coordination |
|  |  |  | Z<sub>A-B</sub> | Lysolipid<sub>1A, 1B</sub> | H-bond |
| Thr188 | Phe<sup>33</sup> | Reduced | Chamber<sub>1-2</sub> | Lysolipid<sub>1A, 1B, 2B</sub> | Lipid tail interaction |
| Phe298 | Ala<sup>33</sup> | Reduced | Chamber<sub>1</sub> | Lysolipid<sub>1A, 1B</sub> | Lipid tail interaction |
| Leu308 | Trp<sup>22</sup> | Reduced | Chamber<sub>1</sub> | Lysolipid<sub>1A, 1B</sub> | Lipid tail interaction |
| Glu309 | Ala/Asp/Arg<sup>22</sup> | Abolished | Z<sub>A</sub> | Lysolipid<sub>1A</sub> | Na<sup>+</sup> coordination |
| Phe312 | Ala/Tyr<sup>22</sup> | Reduced | Chamber<sub>1</sub> | Lysolipid<sub>1A, 1B</sub> | Lipid tail interaction |
| Ile333 | Ala<sup>22</sup> | Reduced | Chamber<sub>1</sub> | Lysolipid<sub>1A, 1B</sub> | Lipid tail interaction |
| Ile341 | Trp<sup>22</sup> | Abolished | Chamber<sub>2-3</sub> | Lysolipid<sub>2B, 3C</sub> | Lipid tail interaction |
| Gln345 | Trp<sup>22</sup> | Reduced | Z<sub>C</sub> | Lysolipid<sub>3C</sub> | H-bond |
| Ala388 | Trp<sup>22</sup> | Reduced | Chamber<sub>1</sub> | Lysolipid<sub>1A, 1B</sub> | Lipid tail interaction |
| Met334 | Ala<sup>22</sup> | Reduced | Chamber<sub>1-2</sub> | Lysolipid<sub>1A, 1B, 2B</sub> | Lipid tail interaction |
| Phe396 | Ala/Ile/Trp<sup>22</sup> | Reduced | Chamber<sub>1-2</sub> | Lysolipid<sub>1A, 1B, 2B</sub> | Gating, Lipid tail interaction |
| Trp400 | Ala/Phe<sup>22</sup> | Reduced | Chamber<sub>1-3</sub> | Lysolipid<sub>1-3</sub> | Gating, Lipid tail interaction |
| Glu421 | Arg<sup>38</sup> | Reduced | Z<sub>A</sub> | Lysolipid<sub>1B, 2B, 3C</sub> | Salt-bridge |
| Thr432 | Ala<sup>33</sup> | Increased | Z<sub>A-B</sub> | Lysolipid<sub>1A</sub> | H-bond |
| Lys433 | Ala<sup>33</sup>, Gln<sup>22</sup> | Abolished/Reduced | Z<sub>A</sub> | Lysolipid<sub>1A</sub> | Salt-bridge |

Like other MFSs, drMfsd2a is comprised of twelve TMs that pack against one another with the substrate binding channel located at a cleft between the N- and C-terminal domains (Fig. 1b-e). The twelve TMs of drMfsd2a are, likewise, subdivided into two helical bundles called the N- (helices 1-6) and C-domains (helices 7-12) that are related by a twofold pseudosymmetry (Fig. 1b). This topology is consistent with other MFSs that have been structurally described, including the recent chicken^22^ and mouse^33^ Mfsd2a structures. These two Mfsd2a structures were solved in the outward-facing, ligand-free conformation from mouse^33^ and the inward-facing state also bound with ALA-LPC from chicken^22^. These two structures are distinct from the conformations reported in this study (Fig. S5). Here, we report five structures of drMfsd2a in the inward-facing conformation ligand-free or with ALA-LPC observed at four distinct locations in the substrate translocation tunnel. The placement of the four ALA-LPC in drMFSD2A is unique from where this lysolipid bound in chicken Mfsd2a ^22^(Fig. 1, S5). In drMfsd2a, the four lysolipids are positioned along the vertical axis of the substrate tunnel within a cleft between the N- and C-domains comprise of helices 1, 2, 4, 5, 7, 8, 9, 10 and 11 that opens to the cytoplasmic side (Fig. 1b-e, S4). Together, the four ALA-LPC binding positions provide a unique opportunity to detail the transport and flippase mechanism of Mfsd2a as the substrate is shuttled through the transporter. Hereafter, the four ALA-LPC captured in drMfsd2a are referred to as Lysolipid_1A,_ Lysolipid_1B_, Lysolipid_2B,_ and Lysolipid_3C_ (number and letter refer the lipid and LPC binding positions) (Fig. 1c-e). Since Mfsd2a transport lipid-LPC from the outer to the inner leaflet, we propose the following order for the four ALA-LPC during substrate transport through Mfsd2a: Lysolipid_1A,_ Lysolipid_1B_, Lysolipid_2B,_ to Lysolipid_3C_.

### Rocker-switch motion

To gain further insights into the differences between two major conformations of the rocker-switch, we mapped the interdomain interactions observed in the outward-facing mouse^33^ and our inward-drMfsd2a structures (Video. S1). The interdomain interactions in the mouse and drMfsd2a structures include hydrophobic, salt-bridge, and hydrogen bond contacts located on the (1) extracellular end, (2) cytoplasmic side, (3) between helices 5 and 8 and (4) across helices 2 and 11 of Mfsd2a (Video. S1). Using the Morphing function in PyMol we calculated 200 states between the transition from the outward mouse^33^ to inward drMfsd2a conformation. This generated a moving model where we observe that the rocker-switch motion in Mfsd2a results from the breaking and forming interdomain interactions mapped to the mouse and drMfsd2a cryo-EM structures. Strikingly, the rocker-switch motion “rocks” along the vertical axes of TM5 and TM8, and TM2 and TM11 (Video. S1, S2a). We also observe that the outward-facing mouse conformation^33^ is held in place by more hydrogen bonds and salt-bridges placed at the opposite, cytoplasmic side than observed for interactions at the extracellular end that locks drMfsd2a in the inward-facing state (Video.S1).

### Rotation of the LPC headgroup

During the transport cycle as lipids traverse from the outer to the inner leaflet, the substrates must translocate and also invert. As such, it is expected that both the acyl tail and headgroup of DHA-LPC will rotate during lipid flipping and translocation through Mfsd2a. Tracing the relative positions between the Lysolipid_1A_ and Lysolipid_1B_ illustrate a significant rotation of the LPC headgroup (Fig. 1d-e, 2). Unlike previous studies that suggest a linear substrate tunnel, our drMfsd2a structures revealed three distinct compartments that each comprise a distinct hydrophobic pocket and a charged cavity to house the lipid tail and LPC headgroup (Fig. 2-3). We term the three hydrophobic pockets Chambers_1-3_ and the three charged cavities the zwitterionic traps_A-C_ (Z_A-C_). Lysolipid_1A_, with the lipid tail in Chamber_1_ and LPC in Z_A_, is the closest substrate to the extracellular side and, hence, likely the first of the four ligand states captured during the substrate transport in the drMfsd2a structures (Fig. 1e, 2a-b). Chamber_1_, also observed in the chicken Mfsd2a structure^22^ but is more extensive in drMfsd2a (Fig. 2a-b, Supplementary text). All the residues in Chamber_1_ are highly conserved (Fig. S4) and mutations at F59, M181, L308, F312, I333, F396, and W400 reduce lysolipid transport activity by Mfsd2a ^22,33^ (Table 1). Based on their placement in Chamber_1_ and Z_A_, the lipid tail and headgroup of Lysolipid_1A_ of are both pointed outwards, bent at an approximately 70° angle (Fig. 1c-e, 2a-b).

**Fig. 2.**
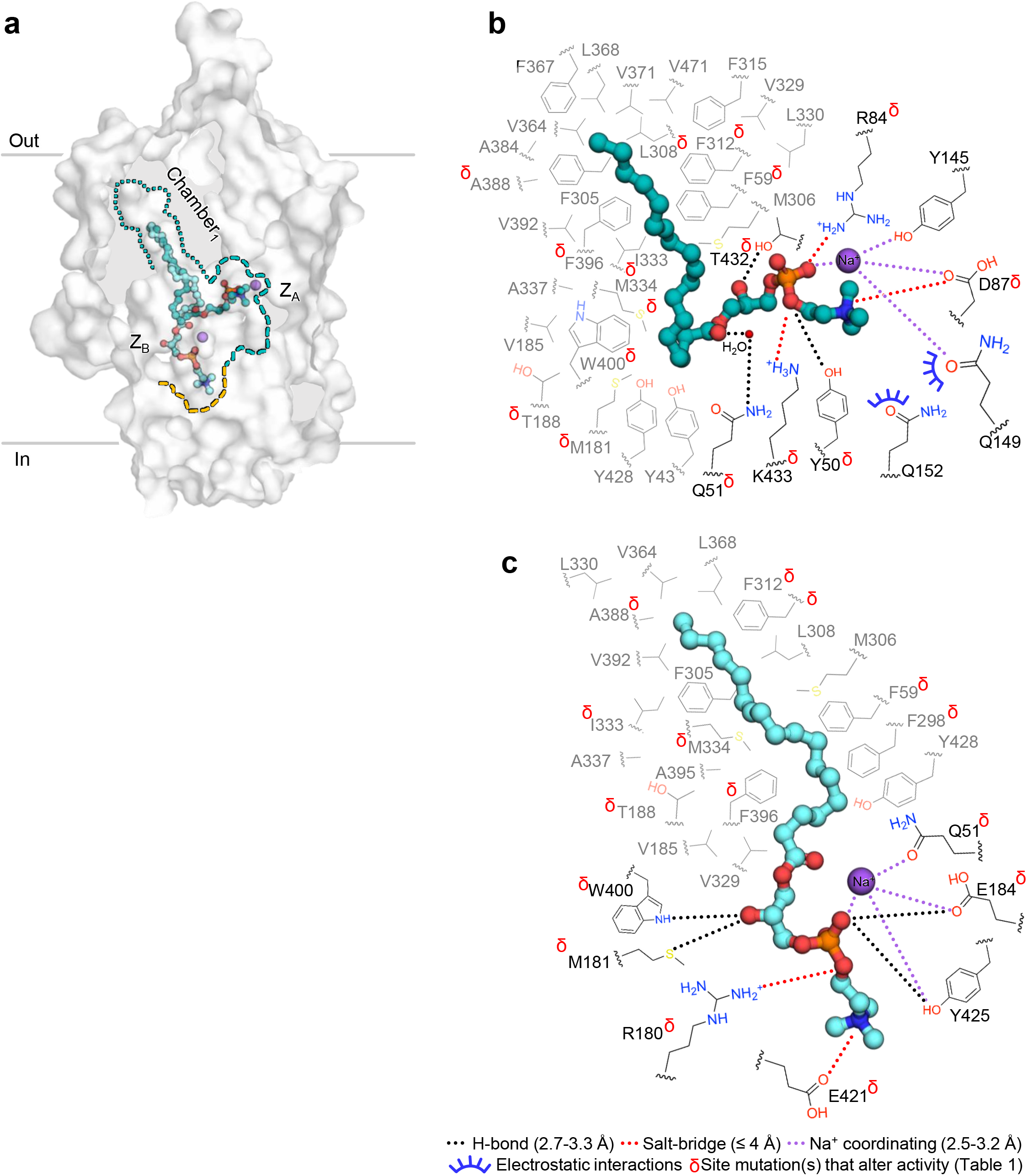
Rotation of the LPC headgroup. **a**, Relative positions of Lysolipid_1A-1B_ in Chamber_1_ and Z_A-B_. Lipid Chamber_1_ is outlined in cyan dotted lines. Z_A-B_ are in dashed cyan and orange lines, respectively. Protein rendered as surface. Lysolipids_1A-1B_ are in stick and sphere. Na^+^ is in purple spheres. **b**, Interactions between lipid tail and LPC of Lysolipid_1A_ with residues in Chamber_1_ and Z_A_. **c**, Interactions between lipid tail and LPC of Lysolipid_1B_ with residues in Chamber_1_ and Z_B_. **b-c**, Carbons of lipid chamber residues are in grey. Carbons of Z-site and Na^+^-ligating residues are in black. Black dotted lines represent H-bonding between 2.6-3.3 Å. Red dotted lines indicate salt bridges with distances ≤ 4 Å. Purple dotted lines are Na^+^ coordinating functional groups within 2.5-3.2 Å. Blue half circles indicate choline coordinating residues within 3.5 Å. Red δ indicate residues with mutations that alter activity. Water molecule shown as small red sphere.

Previous studies established that Mfsd2a transports lipid-LPC in a Na^+^ dependent manner^6,10^. It is proposed that the Mfsd2a symporter uses the flow of Na^+^ down its electrochemical gradient, from the out to inside the cell, to drive the conformational changes for the transport and flipping of lysolipids^6,10^. Molecular dynamics and functional analysis of the chicken Mfsd2a proposed conserved and dynamic Na^+^ binding sites annotated as D87, D91, E309, and K433 (drMfsd2a sequence numbers)^22^. These studies, however, do not provide clear structural details about how Na^+^ orchestrates lysolipid transport and flipping by Mfsd2a and require further investigation. In the Lysolipid_1A_ structure, we observe a nearby unmodeled density that cannot be attributed to the protein or the nearby ALA-LPC Lysolipid_1A_ (Fig. S6a). When a Na^+^ is placed, this cation can be coordinated in a planar molecular geometry by D87, Q149, and Y145 but instead of an amino acid, the fourth site is directly ligated to the phosphate of LPC of Lysolipid_1A_ (Fig. 2b, S6a). This type of planar ligation geometry is a common feature observed for other Na^+^ binding proteins^36,37^. We refer to this cation as Na_A_. In this configuration, the zwitterion phosphate and choline groups of Lysolipid_1A_ is rigidified by (1) salt bridges to D87, R84 and K433, (2) hydrogen bonds to Y50, Q51, and T432, and (3) electrostatic interactions with Q149 and Q152 (Fig. 2b, S6a). This LPC binding arrangement is supported by studies illustrating that Mfsd2a can only transport lysolipids with headgroups that comprise of (1) a negative charge, but not necessarily a phosphate, and (2) amine moiety^30,35,38^. We term the Mfsd2a interactions with the Lysolipid_1A_ LPC as zwitterion trap_A_ (Z_A_). Consistent with these observations, Y50, Q51, D87, R84, T432, and K433 are highly conserved (Fig. S4) and their mutations decrease or prohibit Mfsd2a lysolipid transport^22,33^ (Table 1). Moreover, molecular dynamics studies from chicken Mfsd2a support the findings that D87 is a primary site for Na^+^ binding^18^.

In the second ligand binding configuration, Lysolipid_1B_, we observed a lipid tail that is also bound in Chamber_1_ but the LPC headgroup is now translocated by 16.9 Å, rotated by ∼70° inwards, bound by a second zwitterion trap, Z_B_ (Fig. 1e, 2a, c, S6b). The Z_B_ site that coordinates the LPC of Lysolipid_1B_ is comprised of (1) two salt bridges by R180 and E421 to the phosphate and choline groups and (2) is further rigidified by hydrogen bonding to M181, E184, W400, and Y425 (Fig. 2c, S6b). This LPC binding arrangement in Z_B_ is, again, support the necessity of a headgroup that contains a negatively charged moiety and amine group for Mfsd2a transport^30,35,38^. Further examination of the Z_B_ site also revealed an unmodeled density adjacent to the phosphate of the LPC headgroup (Fig.2a, c, S6b). When modeled as a Na^+^, this cation can be coordinated in a planar molecular geometry by Q51, E184, Y425, and the phosphate of the LPC of Lysolipid_1B_ (Fig. 2c, S6b). We refer to this Na^+^ as Na_B_. All residues that comprise Z_B_ and the Na_B_ coordination sites are conserved (Fig. S4) and mutations of Q51, M181, R180, E184, W400, and E421 reduce Mfsd2a lipid-LPC transport activity^22,33,38^ (Table 1). Unlike the LPC headgroup, the translocation of C18 of the Lysolipid_1B_ lipid tail from Lysolipid_1A_ is more modest and still adhered in Chamber_1_ (Fig. 1a, 2a, c). Due to the shifts in the Lysolipid_1B_, Chamber_1_ is modified for binding the lipid tail with fewer interactions (Fig. 2c, Supplementary text). Together, the trajectory from Lysolipid_1A_ to Lysolipid_1B_ describes a ∼70° rotation and 16.9 Å translocation of the LPC from a head-out to a head-in orientation (Fig. 1d-e, 2, S6a-b).

### Lipid translocation

After the rotation of the LPC headgroup, we propose that the next steps during lysolipid transport involve the translocation of the substrate by Mfsd2a towards the cytoplasmic side (Fig. 3). In the Lysolipid_2B_ configuration, the C18 of the ALA lipid tail is displaced 13.9 Å relative to Lysolipid_1B_ (Fig. 1a, c, e) and is bound in Chamber_2_ located towards the cytoplasmic side of drMfsd2a (Fig. 3a-b). Chamber_2_ is shifted inward from Chamber_1_ and is comprised of highly conserved residues (Fig. 3b, S4, Supplementary text) and mutation of M181, T188, and I341 disrupt or abolish Mfsd2a transport activity^22,33^ (Table 1). Unlike the large movement in the lipid tail, the LPC of Lysolipid_2B_ is bound in a similar position to Lysolipid_1B_ by Z_B_. The phosphate and choline of Lysolipid_2B_ are also held in place by salt bridges to R180 and E421 but is now shifted more towards the cytoplasmic side, forming hydrogen bonds with H156 (via a water), S160, and E184 (Fig. 3a-b, S6c). Instead of the phosphate and a fourth amino acid like Lysolipid_1A_ and Lysolipid_1B_, the hydroxyl moiety of the headgroup and a water forms two of the ligation sites to Na^+^ along with Q51 and E184. Consistent with our findings, Q51, S160, H156, R180, E184, and E421 are conserved (Fig. S4). Mutation of Q51, S160, H156, R180, E184, and E421 disrupt or abolish Mfsd2a transport activity^19,22,33^ (Table 1). It is noteworthy that mutation of S160 is associated with decreased DHA transport to the brain and results in severe microcephaly^13^.

**Fig. 3.**
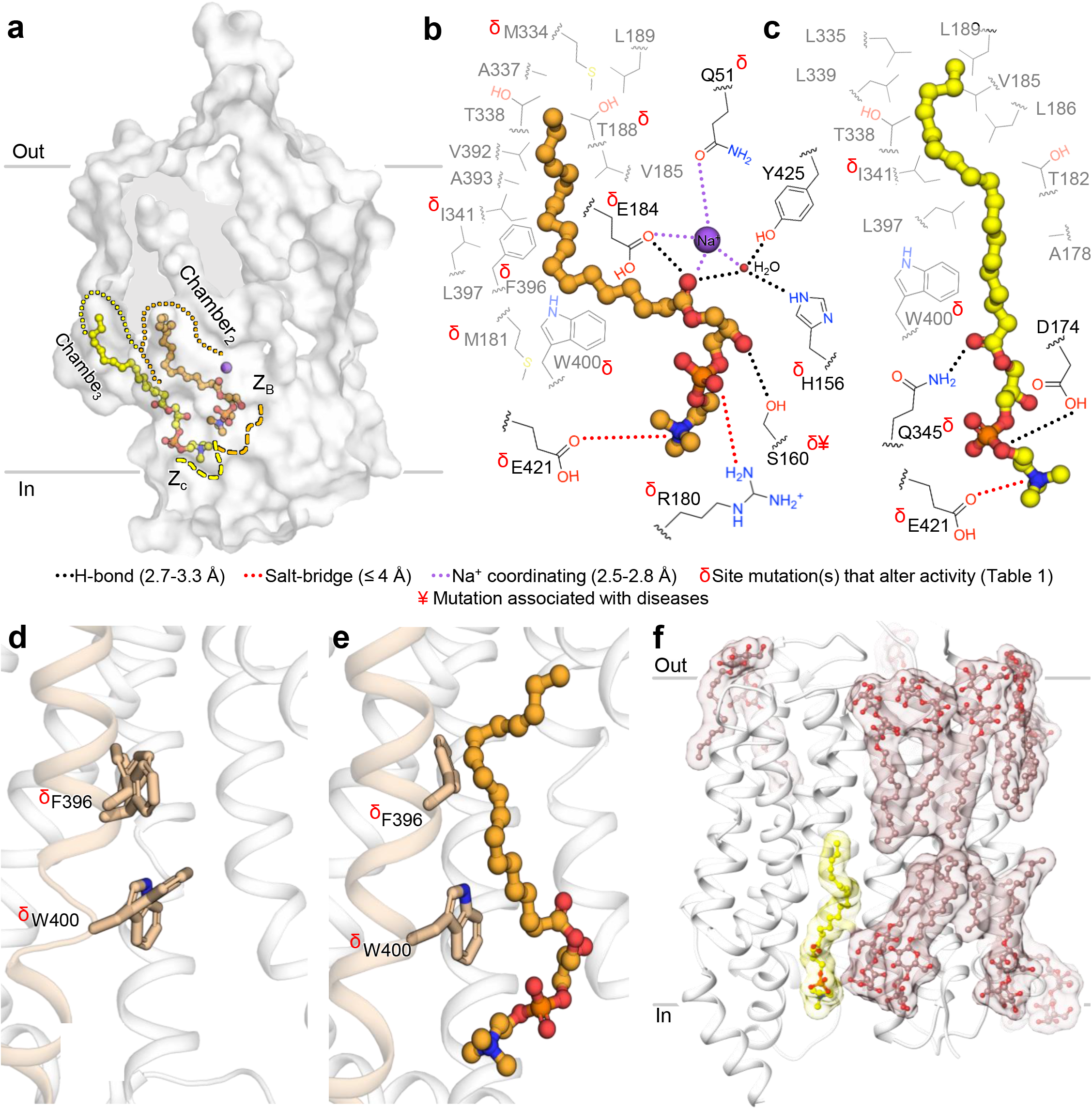
Lysolipid translocation and release through Chamber_2-3_ and Z_B-C_. **a**, Relative positions of Lysolipid_2B-3C_ in Chamber_2-3_ and Z_B-C_. Lipid Chambers_2-3_ are outlined in orange and yellow dotted lines, respectively. Z_B-C_ are in dashed orange and yellow lines, respectively. Protein is rendered as surface. Lysolipids are in stick and sphere. Na^+^ is in purple spheres. **b**, Interactions between lipid tail and LPC of Lysolipid_2B_ with residues in Chamber_2_ and Z_B_. **c**, Interactions between lipid tail and LPC of Lysolipid_3C_ with residues in Chamber_3_ and Z_c_. **b-c**, Lipid chamber residues are in grey. Z-site and Na^+^-ligating residues are in black. Black dotted lines represent H-bonding between 2.6-3.3 Å. Red dotted lines indicate salt bridges with distances ≤4 Å. Purple dotted lines are Na^+^ coordinating functional groups within 2.5-2.8 Å. Red δ and ¥ indicate residues with mutations that alter activity and are associated with disease (see Table 1). Water molecule shown as small red sphere. **d**, Substrate transport tunnel in closed conformation. The Lysolipid_1A_ structure is shown with alternate conformations of the F396 and W400 from structures without lipid tail of ALA-LPC in Chamber_2_. **e**, Substrate transport tunnel in open conformation with Lysolipid_2B_ bound. **d-e**, Protein rendered as cartoon. Gating residues are in stick. TM10 colored in wheat. **f**, Lysolipid_3C_ (yellow) at the cytoplasmic exit shown adjacent to the DDM belt with detergent molecules shown in stick and ball, and in tan transparent surface.

Examination of Lysolipid_3C_ revealed that the ALA-LPC lipid tail is in a third hydrophobic chamber (Chamber_3_) located adjacent to the cytoplasmic opening of the drMfsd2a substrate tunnel (Fig. 3a, c). Comparison with Lysolipid_2B_ indicates that the C18 of the lipid tail and choline of Lysolipid_3C_ have translocated inwards by 8.1 Å and 4.1 Å (Fig. 1c-e). The more modest sized Chamber_3_ is exposed to the exterior of the cytoplasmic side of drMfsd2a (Fig. 1e, 3a, c, Supplementary text). All residues that comprise Chamber_3_ are highly conserved (Fig. S4, Supplementary text) and mutation of I341 and W400 limit or abolish Mfsd2a transport activity^22^ (Table 1). Similar to the fewer interactions that bind the lipid tail, the LPC of Lysolipid_3C_ is coordinated by a single salt bridge of the choline by E421 and two hydrogen bonds to D174 and Q345 (Fig. 3c, S6d). Consistent with these observations, D174, Q345 andE421 are highly conserved (Fig. S4) and mutation of the latter two decrease or prohibit Mfsd2a lysolipid transport activity^22,38^ (Table 1). Together, the position from Lysolipid_1B_ to Lysolipid_3C_ describes a 22 Å and 9.4 Å translocation of the C18 of the lipid tail and the choline of the headgroup (Fig. 1c-e).

### Open and closed conformations of the Mfsd2a gates

In addition to the residues that make up the three hydrophobic chambers, it has been proposed that substrate “gating” of Mfsd2a is comprised of bulky hydrophobic residues that respond to the presence of and orchestrate the conformational changes required for transport of lysolipids^39^. Molecular dynamic simulation of the chicken Mfsd2a proposed inner gates that are located within the hydrophobic tunnel consisting of the conserved residues M181, F396, and W400^22^ (Fig. S4). Indeed, we observe movements of F396 and W400 but not M181 that is dependent on the presence of a nearby acyl-chain in drMfsd2a (Fig. 3d-e). Here, F396 and W400 interact with lipid tails in Chamber_1-2_ and Chamber_1-3_, respectively (Fig. 2-3). In the structure with the lipid tail absent from Chamber_2_, F496 and W400 are flexible causing a break in TM10 (Fig. 3d). Here, F396 and W400 occupy alternate conformations that block access to Chamber_2_. In the presence of lipid tail of Lysolipid_2B_ in the Chamber_2_, F396 and W400 are rigidified to reform the disordered portion of TM10 (Fig. 3e). In this structure, F396 and W400 are flipped to a single conformation allowing access to Chamber_2_. F396 and W400 are now also engaged in forming Van der Waal interactions with the acyl-chain of the Lysolipid_2B_ within Chamber_2_ (Fig. 3e). We propose that the “gating” interactions with these bulky hydrophobic residues facilitate the translocation of the lipid tails from Chamber_1_ through Chamber_3_. Consistent with our the above postulate, functional analyses coupled with mutagenesis of F396 and W400 reduce Mfsd2a substrate transport^22^ (Table 1). Moreover, molecular dynamics simulation indicates similar findings where the presence of Na^+^ and lipid cargos stimulate the opening of the F396 and W400 gates^22^ to allow lipid tail access to Chamber_2_.

### Lipid and Na^+^ release

Based on its unique position, the following features observed in our studies suggest Lysolipid_3C_ is poised for release. First, there are fewer contacts between the lipid tail and hydrophobic residues of Chamber_3_ than Chamber_1-2_ (Fig. 2-3). Second, there are also minimal contacts between LPC of Lysolipid_3C_ and Z_C_ (Fig. 3c, S6d). Third, Lysolipid_3C_ is placed in a groove that is exposed on the cytoplasmic side of drMfsd2a (Fig. 1c-e, 3a, c, 3f). Fourth, when mapped to the adjacent DDM molecules, Lysolipid_3C_ is closely aligned with the detergent belt that encircles the cytoplasmic side of drMfsd2a (Fig. 3f). Given these rationales, we propose that Lysolipid_3C_ is poised for release to through the cytoplasmic opening of the substrate tunnel of Mfsd2a for integration into the inner membrane leaflet (Fig. 1e, 3f). Further, we do not observe density for Na^+^ near Lysolipid_3C_. The absence of a density that can be attributed to Na^+^ at this site may be due to the current resolution of our studies and/or intrinsic dynamics of the exterior protein region. It is also possible that the absence of a Na^+^ density indicates that cation was released during the transition of Lysolipid_2C_ to Lysollipid_3C_ (Fig. 3c, 4f-h, S6).

### Proposed Lipid flipping and Mfsd2a cycling

During the transport of the substrate, Mfsd2a performs the energetically unfavorable task of flipping the lysolipid from the outer with the inner membrane leaflet^6,28^ (Fig. 4). To our knowledge, a molecular model of the lipid flipping mechanism has not been structurally detailed for flippases or floppases. The only other protein with structural mechanism for lipid flipping is the recently determined structure of the TMEM16 scramblase^29^. However, unlike Mfsd2a, TMEM16 does not rely on coordinated interactions between protein and lipids cargos. Instead, TMEM16 lowers the activation energy for lipid transport and flipping by thinning the immediate surrounding membrane^29^. Using the five structures determined here, together with the previously determined structures of Mfsd2a, we propose that lipid flipping occurs in a series of steps by which the lipid head and tail are interchangeably anchored to the transporter as the lipid traverses through the substrate tunnel.

**Fig. 4.**
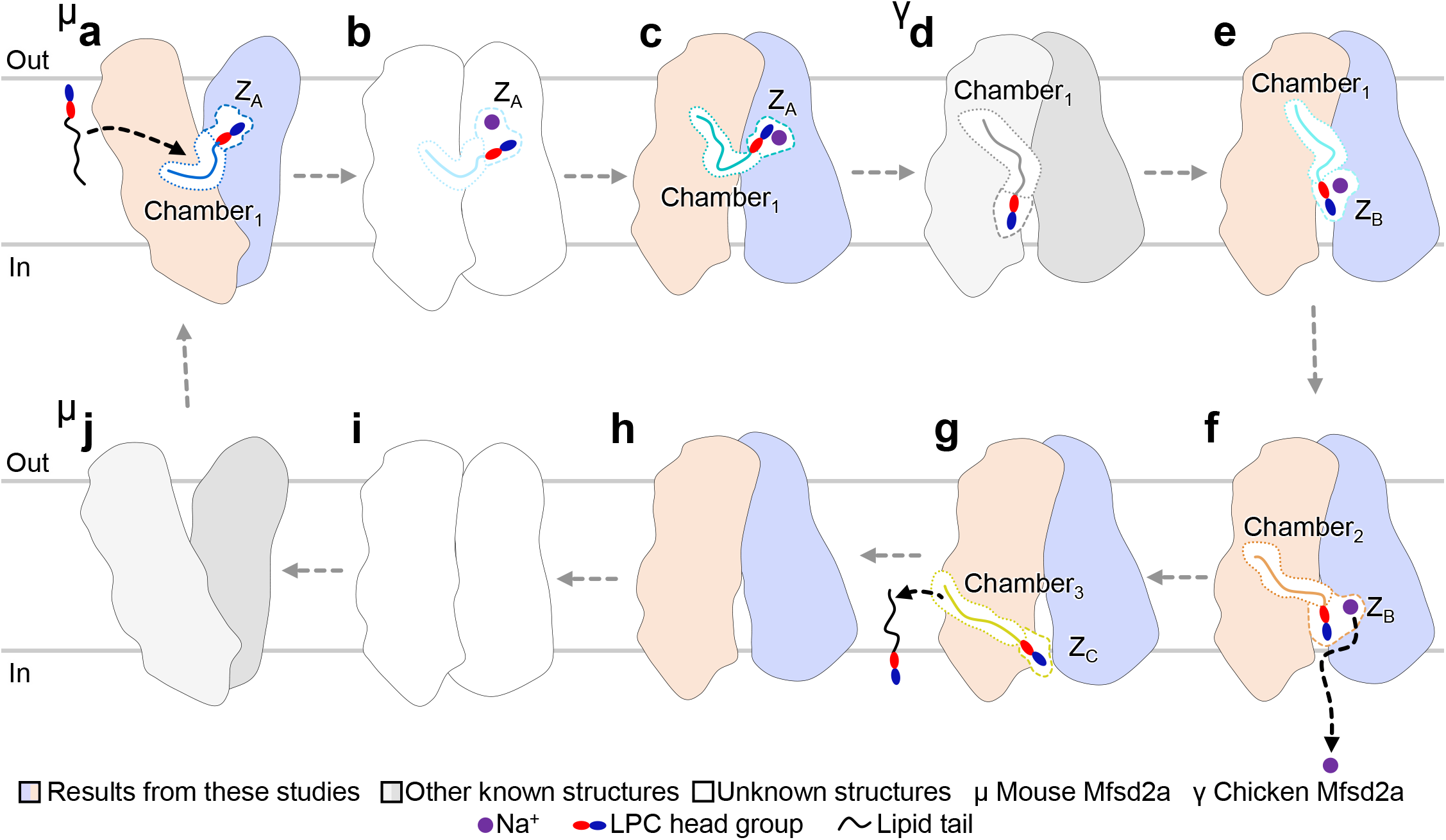
Proposed model for Mfsd2a lipid transport cycle. **a**, The ligand-bound, outward-facing mouse Mfsd2a structure docked with ALA-LPC is bound in a lateral position Chamber_1_ (see Supplementary text). The LPC headgroup is trapped by Z_A_ in an outward pointing orientation. **b**, The proposed and unknown occluded ligand-bound conformation. Mfsd2a rocks from an outward to inward-facing conformation. **c-e**, The various conformations during flipping of the LPC headgroup. The lipid tail is held in position by Chamber_1_ while the LPC headgroup samples multiple configurations between Z_A_ and Z_B_ before it is moved to Z_B_. This movement flips the LPC headgroup from an outward to inward-pointing orientation. This process results in a substrate with the lipid tail pointing out and the LPC headgroup pointing in, a configuration that is aligned with the inner membrane leaflet. **f**, The translocation of the lysolipid to Chamber_2_ and Z_B_. The lipid tail is translocated from Chamber_1_ to Chamber_2_. The LPC headgroup is shifted inwards. This process translocates the entire lipid-LPC substrate closer to the cytoplasmic exit. **g**, The release of the substrate to the cytoplasmic side. LPC headgroup is moved to Z_C_ and the lipid tail is translocated to Chamber_3_. Chamber_3_ and Z_C_ are located at the cytoplasmic exit. **h**, The substrate and Na^+^-free structure of drMfsd2a in the inward-facing conformation. This structure represents the conformation after lysolipid release from Mfsd2a. **i-j**, The conformation during resetting of the transporter. In the absence of the substrate, Mfsd2a resets to the outward-open conformation (j) by first transitioning through a ligand-free occluded state (i).

Since the lipid translocates from the outer to the inner membrane leaflet, the first anchoring event observed is the transition between Lysolipid_1A_ to Lysolipid_1B_ where the lipid tail is attached to Chamber_1_ while the head group is free to flip into from Z_A_ to Z_B_ (Figure 2, 4c-e, Video S2b). Once the headgroup is attached to Z_B_, as observed for Lysolipid_1B_ (Fig. 2c), the tail is released from Chamber_1_ and anchored to Chamber_2_ where Lysolipid_2B_ is found (Fig. 3b). While the four lipid-LPC span the length of the drMfsd2a substrate tunnel (Fig. 1e), key steps are still missing. Namely, it is unclear whether the lipid tail or LPC anchors first to Chamber_3_ or Z_3_ (Figure 3). Our structures do demonstrate that the translocation of the entire lipid-LPC occurs through an exit in the substrate tunnel that opens to the cytoplasmic side (Figure 1e, 3, 4f-g, Video S2c).

Moreover, we would like to suggest a model for the co-transport of Na^+^ and the lipid through Mfsd2a based on the proposed cation binding in Lysolipid_1A_, Lysolipid_1B_, and Lysolipid_2B_. These structures suggest direct ligation between the Na^+^ and ALA-LPC (Fig. 2-3, S6). If true, it is tempting to suggest a model of the coupled movement of Na^+^ with lipid-LPC down the electrochemical gradient of the cation during translocation and flipping of the lysolipid through Mfsd2a (Fig. 2-3, 4c-g, Video S2). Together, the lipid-tail versus LPC anchoring model provides a reasonable mechanistic process where the lipid flips and translocates in a stepwise manner through the chambers until it emerges out at the apposing end and inserts into the inner membrane leaflet. While more remains to be determined in future studies, a detailed proposal of the Na^+^-dependent steps of lipid flipping and translocation through Mfsd2a can be found in the Supplementary text.

### Concluding remarks

In this study, five structures of drMfsd2a were described and four unique lipid-LPC binding configurations identified. The structures allow us to propose a Na^+^-dependent, stepwise lipid flipping mechanism of Mfsd2a. Extensive mutagenesis and functional analyses confirm the importance of key residues identified for lysolipid transport and flipping. Moreover, we identify two Na^+^ binding sites, including Na_B_ which has not been observed previously, and describe the possible role that this cation plays in omega-3 fatty-acid binding, flipping, and release. While these snapshots allow us to propose a stepwise lipid flipping mechanism, there remains several unanswered questions. For example, is the sodium required for the stepwise translocation of the lipid or simply to stabilize the headgroup? When does acyl-chain inversion occur? Here, we have preliminary docking simulations that suggest that acyl-chain inversion happens before the rotation of the headgroup (Fig S7 versus 1e). What role does protein conformation change from outward to inward play in this process? Is the protein simply acting as a scaffold for lipid flipping or is the rocker-switch critical in ushering the lipid through this process? Future studies must delve more deeply into the proposed flippase and lipid transport for Mfsd2a and for other less common lipids that play critical roles in neuromodulation and viral pathogenic pathways.

## Methods

### Protein expression and purification

Mfsd2a isoform B from zebrafish (Uniprot Q6DEJ6) was subcloned in the pFasbac1 vector (Invitrogen) with a N-terminal-10Xhis tag. The Mfsd2a sequence was truncated at residue D22 of the N-terminus, D509 of the C-terminus, and with the N509Q, N214Q, N225Q mutations to remove post-translational glycosylation modifications. DH10bac was used for preparation of bacmids. Recombinant baculovirus was generated using the *Spodoptera frugiperda* Sf-9 system and transfection following the protocol provided for the Bac-to-bac Baculovirus Expression System. Mfsd2a -pFasbac1 construct was used to transfect into Sf-9 cells following standard protocol to generate three generations of baculovirus. P3 generation of the Mfsd2a -pFasbac1 baculovirus was use to infect batch culture of Sf-9 for protein expression.

Mfsd2a was overexpressed in Sf-9 insect cells, harvested 60 h after infection. The cell pellets were resuspended in lysis buffer containing 20 mM Tris (pH 8.0) and 150 mM NaCl and supplemented with EDTA-free protease inhibitor cocktail (Roche). The Sf-9 cells were lysed using 50 homogenizing cycles on ice. The lysate was clarified by centrifugation at 130,000 g for 1 h. The pelleted membrane was resuspended in high-salt buffer containing 1.6 M NaCl and 20 mM Tris (pH 8.0) for washing and centrifuged again for 1 h at 130,000 g to remove soluble debris. The pelleted membrane was rapidly frozen in liquid nitrogen and stored at –80°C until further use.

To purify Mfsd2a, the membrane pellet was solubilized and membrane protein extracted in 2% n-dodecyl-β-d-maltopyranoside (DDM), 20 mM Tris (pH 8.0), 150 mM NaCl, 5% glycerol and 0.2% cholesteryl hemisuccinate Tris salt (CHS) for 4 h at 4°C. After extraction, the insoluble debris were removed by centrifugation for 1 h at 130,000 g. 20 mM imidazole (pH 8.2) was added to the supernatant and incubated with TALON beads for 16 h at 4°C. The Mfsd2a -bound beads were washed with 6 column volumes of 20 mM imidazole, 20 mM Tris (pH 8.0), 500 mM NaCl and 0.1% DDM. The resin was then equilibrated in buffer composed of 20 mM Tris (pH 8.0), 150 mM NaCl, 0.4% decyl-β-d-maltoside (DM) and 0.02% DDM. At 4°C, the 10 × His tag was removed by on-column thrombin digestion overnight at an enzyme:protein molar ratio of 1:500. The cleaved Mfsd2a was collected in the flow-through and was flash frozen in liquid nitrogen and stored at – 80°C until further use. FAB production and purification

Mouse IgG monoclonal antibodies against drMfsd2a were produced by the Monoclonal Antibody Core (D. Cawley). 330 µg of purified Mfsd2a in buffer containing 20 mM Tris (pH 8.0), 150 mM NaCl, 0.02% DDM and 0.002% CHS was used to immunize mice in three injections. 15 × 96-well plate fusions yielded 169 IgG-positive wells at a 1:30 dilution. Native and denatured Mfsd2a proteins were then used in ELISA to search for candidates that bound the conformational epitopes27, where Ni-NTA plates were used for Mfsd2a immobilization. Thirty-five of the 169 fusions showed significant preference for binding against well-folded Mfsd2a. Western blotting was performed to assess the binding affinity and specificity of the antibodies generated from hybridoma cell lines. Monoclonal antibody 11D3 was then purified from the hybridoma supernatants by 4-mercaptoethylpyridine (MEP) chromatography. FAB fragments were produced by papain digestion and purified in flow-through buffer containing 20 mM NaPi (pH 8.0) and 150 mM NaCl by protein A affinity chromatography.

### Mfsd2a-FAB complex assembly

Purified 15A9 FAB fragment was incubated overnight at 4°C with Mfsd2a at a 5:1 molar ratio. DrMsfd2a–15A9 complex was injected into the size exclusion column Superdex200 to separate the unbound FAB. The size exclusion column step was also used to exchange the buffer of the drMfsd2a -FAB complex into 50 mM HEPES (pH 8.0), 150 mM NaCl and 0.02% DDM. SDS-PAGE analysis was used to pool peak fractions containing both dr Mfsd2a and FAB, indicating complex formation (Fig.S1). The pooled fractions containing Mfsd2a -FAB were pooled and concentrated to 3 mg/ml using Amicon spin concentrator with a 30 kDa cutoff.

### Mfsd2a-FAB Cryo-EM grid preparation

400-mesh 1.2/1.3 Cu or 300-mesh UltrAuFoil 1.2/1.3 Au grids (Quantifoil) were made hydrophilic by glow discharging for two times 60 seconds with a current of 15 mA in a PELCO easiGlow system. The cryo grids were prepared using a Leica EM GP (Leica). The chamber was kept at 4°C and 95% humidity (86-91% measured). 3 µl sample at 3 mg/ml were applied to a glow-discharged holey grids, blotted for 6 s, and plunge frozen into liquid ethane and stored in liquid nitrogen.

### Mfsd2a -FAB cryo-EM data collection

Cryo-EM grids were loaded into a 300 keV FEI Titan Krios cryo-electron microscope (ThermoFisher Scientific, formerly FEI) at HHMI Janelia Reasearch Campus, Janelia Krios 1, equipped with a C_s_ corrector, and Gatan energy filter and K3 camera (Gatan Inc.). Movies of 50 frames with 1 e^-^/Å^2^ per frame (50 e^-^/Å^2^ total dose) were automatically recorded at a nominal magnification of 81,000x, corresponding to a physical pixel size of 0.844 Å/px (superresolution pixel size 0.422 Å/px) in CDS mode at a dose rate of 9.5 e^-^/px/s (∼7.5 e^-^/px/s on the camera through the sample) and a defocus range of -0.5 to -1.8 µm using SerialEM^40^. In total, 2,653 and 8,443 movies were collected in two separate imaging sessions, with the first dataset on a 400-mesh copper grid and the second on a 300-mesh UltrAuFoil gold grid (Table S1).

### Mfsd2a-FAB cryo-EM data processing

The overall workflow of the image processing is illustrated in Fig. 2. All preliminary steps were performed within RELION 3.1^41^ unless specified. Movie alignment and dose weighting were performed with MotionCor2^42^ in a 5 × 5 patch. Contrast transfer function (CTF) parameters were estimated with Gctf^43^. Good 2D classes generated from ∼1,000 manually picked particles served as templates for automatic particle picking in RELION, resulting in 623,644 and 3,033,688 particles in dataset 1 and dataset 2 respectively. Particle images extracted were 4× binned, resulting in a pixel size of 3.376 Å, and then subjected to several rounds of 2D classification and 3D classification, removing junk particles and particles without FAB bound. Particles were re-extracted, unbinned to a pixel size of 0.844 Å, and subjected to 3D auto-refinement with local angular searches (RELION additional arguments: --sigma_ang 1.667) followed by Bayesian polishing. Polished particles from 2 datasets were imported into cryoSPARC v3.3.2^44^. For dataset 1, 146,914 particles were subjected to CTF refinement to correct for beam-tilt, spherical aberrations, and per-particle defocus parameters, followed by non-uniform refinement resulting in a map at 3.3 Å. For dataset 2, 346,394 particles were first cleaned up by heterogenous refinement, followed by CTF refinement and non-uniform refinement resulting in a map at 3.2 Å. Two datasets were merged and further classified by heterogenous refinement and refined to 2.9 Å with local resolution regions up to 2.4 Å using non-uniform refinement and local resolution filtering. Altogether, 60 maps (from cryoSPARC) between 2.9 to 4.13 Å were created using different strategies of processing the data using cisTEM^45^, RELION^41^, and cryoSPARC^44^, showing different degrees of details, especially around the ligand binding sites.

Densities for three ligands in the 2.9 Å map from 295,580 particles were clearly observed in the cavities of Mfsd2a which are unlikely to co-exist simultaneously. We hypothesized that the three ligand densities shown in that particular density map are a result of merged particle populations where subsets with different ligand positions have not been classified due to the small ligand size. To sort out subsets with different ligand positions, 3D variability analysis^46^ and 3D classification without alignment using a tight mask covering the ligand binding sites were used without obvious success. Alternatively, reference-based 3D classification without alignment was deployed. Individual ligand densities were segmented and extracted with UCSF Chimera^46^. Extracted ligand densities were then subtracted from the Mfsd2a-FAB map, generating references with (1)3 ligands bound, (2) no ligand bound, (3) only ligand 1 bound, (4) only ligand 2 bound, and (5) only ligand 3 bound, respectively. These maps served as references for 3D classification without alignment, with a tight mask covering ligand binding sites included. To avoid reference bias and overfitting, individual particle subsets obtained from 3D classification were subjected to ab-initio reconstruction, followed by non-uniform refinement, resulting in maps at 3.3 Å, 3.4 Å, 3.3 Å, 3.4 Å, and 3.4 Å, with local resolutions to up to 2.8 Å (3-ligands, lower resolution), 2.8 Å (ligand-free), 2.8 Å (1A), 2.9 Å (2B), and 2.9 Å (3C), respectively (Fig. S2-3, Table S1-2). The map with the clearest Lysolipid_1B_ configuration was generated from 413,435 particles using a combination of processing in cisTEM^45^ and cryoSPARC^44^ and has an average resolution of 4.1 Å with up to 2.5 Å locally (Table S2). Six of the >60 maps have been made available and have the following accession codes: EMD-27148, EMD-27149, EMD-27150, EMD-27151, EMD-27152, and EMD-27153.

The initial model fitting was achieved by manual rigid body rotation and translation and the “Fit Map” of the chicken Mfsd2a structure^22^ (PDB 7MJS) into the 2.9 Å electron density maps for drMfsd2ain UCSF Chimera^46^. The protein sequence of the structures was altered to match drMfsd2a and manually fitted into the maps in COOT^47^. Building of the FAB portion of the structure was achieve similar to drMfsd2a starting with the FAB model from the SLC38A9-FAB^48^ structure (PDB 6C08). Refinement of the drMfsd2a structure was accomplished by iterative cycles of manual fitting and automatic refinement in Coot^47^ and Real Space Refine in Phenix^49,50^, respectively. The final structure validation was carried out in Real Space Refine in Phenix^50^ (Table S2). The fitted pdbs have been made available together with the EM maps with the following PDB IDs: 8D2S, 8D2T, 8D2U, 8D2V, 8D2W, and 8D2X. Figures and movies were generated in Pymol or UCSF Chimera^46^.

## abbreviations

(Mfsd2a): MFS domain containing protein-2a
(DHA): docosahexaenoic acid
(ALA): α-Linolenic acid
(LPC): lysophosphatidylcholine
(BBB): blood brain barrier
(MFS): major facilitator superfamily
(CNS): central nervous system
(cryo-EM): cryo-electron microscopy
(TMs): transmembrane helices

## Author contribution

C.N., H.T.L., M.J.G. performed cloning, protein expression, purification, antibody generation and complex formation, and initial screening by cryoEM; C.N., H.T.L., M.J.G., D.M. performed cryoEM screening and data collection; C.N., H.T.L., L.T.F.L., M.J.G., D.M. performed single particle reconstructions, worked on model building and refinement. All authors participated in manuscript preparation, figure preparation, and interpretation of the data and analyses. H.T.L. and T.G. conceived the project.

## Acknowledgements

The authors thank Rui Yan at the HHMI Janelia research campus for her help with data collection. We also thank the Janelia Research Campus cryo-EM center for the use of their electron microscope. This work utilized the computational resources of the NIH HPC Biowulf cluster (http://hpc.nih.gov). The Gonen laboratory is supported by funds from the Howard Hughes Medical Institute and by the National Institutes of Health P41-GM136508.

## Supplementary Information

### Supplementary text

#### Lipid-drMfsd2a interactions

The interactions between drMFS2A and the lipid tails were identified by visual inspection of residues that surround the acyl-chain for each ALA-LPC in COOT. These interactions are listed as below and illustrated in Figures 2-3.

Chamber_1_ - Lysolipid_1A_: Residues Y43, F59, M181, V185, T188, F305, M306, L308, F312, F315, V329, L330, I333, M334, A337, V364, F367, L368, V371, A384, A388, V392, F396, W400, Y428, and V471 (Fig. 2a-b).

Chamber_1_ - Lysolipid_1B_: Residues F59, V185, T188, F298, F305, M306, L308, F312, V329, L330, I333, M334, A337, V364, L368, A388, V392, A395, F396, W400, and Y428 (Figure 2a, c).

Chamber_2_ - Lysolipid_2B_: Residues M181, V185, T188, L189, M334, A337, T338, I341, V392, A393, F396, L397, and W400 (Figure 3a-b).

Chamber_3_ - Lysolipid_3C_: A178, T182, V185, L186, L189, L335, T338, L339, I341, L397, and W400 (Figure 3a, c).

#### A proposed model for lipid flipping and Mfsd2a cycling

During the transport of the substrate, Mfsd2a performs the energetically uphill task of flipping the substrate from a head-out, tail-in to a head-in, tail-out orientation to align the lysolipid from the outer with the inner membrane leaflet^6,28^ (Fig. 4). To our knowledge, a molecular model of the lipid flipping mechanism has not been structurally detailed for flippases or floppases. The only other protein with structural mechanism for lipid flipping is the recently determined structures of the TMEM16 scramblase^29^. However, unlike Mfsd2a, the TMEM16 mechanism does not rely on coordinated interactions between protein and lipids cargos to translocate and flip the substrates. Instead, TMEM16 lowers the activation energy for lipid transport and flipping by thinning the immediate surrounding membrane^29^.

To gain further insights into the initial binding and flipping of the lysolipid, we performed docking experiments of ALA-LPC on the outward-facing mouse Mfsd2a structure^33^ (Fig. S7). There are three key findings in our docking studies. First, the lipid tail is bound in a lateral orientation, dictated by the shape of a modified Chamber_1_ that faces outwards to the extracellular side (Fig. 4a, S7). Second, we observe that the docked LPC is bound in the same cavity that comprise residues of Z_A_ (Fig. 2, 4a, S6a, S7). Third, similar to our Lysolipid_1A_, the LPC is pointed outward, an orientation of the lysolipid that is still aligned with the outer leaflet (Fig. 4a, S7). We believe these docking studies represent a lysolipid binding configuration before the transporter transitions from an outward to an inward-facing conformation, stimulated by ligand and possibly Na^+^ binding (Fig. 4a-c, S6-7). We propose that during this inward rocking motion, Chamber_1_ changes from an outward-facing, lateral position to an inward, vertical conformation (Fig. 2, 4a-c, S7). If true, the change in conformation of Chamber_1_ to a vertical position is likely the driving force to flip the lipid tail from a lateral to a vertical, outward pointing orientation, as observe for Lysolipid_1A_ (Fig. 1e, 2a-b, 4c, Video S2). Moreover, because the headgroup of Lysolipid_1A_ is still in Z_A_ after Mfsd2a changes from the outward to inward-facing conformation, the flipping of the acyl-chain outwards results in a bent Lysolipid_1A_ where both the lipid tail and LPC are pointing outwards (Fig. 1c, 2a-b, 4a-c, Video S2). Given these results, we propose that the inversion of the acyl-chain takes place before the headgroup. Specifically, we propose that the reorientation of the lipid tail to point outwards occurs during the transition of Mfsd2a from the outward to inward-facing conformation and is the first key step in the flipping of the lysolipid to align with the inner membrane leaflet (Fig. 4, Video S2).

After the reorientation of the acyl-chain to point outwards, the largest rotation of the headgroup occurs between the transition from Lysolipid_1A_ to Lysolypid1_B_, a process that we propose is facilitated by Na^+^. Based on the placement of the Na_A-B_ (Fig. 2-4, S6, Video S2), we propose that the transport of Na^+^ plays key role in translocating and flipping of the lysolipid. When LPC is present at Z_A_, the Na^+^ is ligated directly to the phosphate group of Lysolipid_1A_ (Fig. 2b, S6a). The phosphate acting as a ligand for Na^+^ binding would provide the rationale for the coupled movement of the lysolipid with Na^+^ down its concentration gradient toward the cytoplasmic side. In addition to ligation, it is likely that Na^+^ acts as a counterion for the phosphate group of LPC. As a counterion, Na^+^ can neutralize the bulky negatively charged phosphate group to facilitate movement of the LPC headgroup and itself through Mfsd2a. In addition to the charged interactions, the flipping of the LPC from Z_A_ to Z_B_ appears to also be facilitated by the following features. First, there is an open cavity between Z_A_ to Z_B_, allowing delocalized LPC movements between the two sites (Fig. 2a). This is consistent with our observation of weaker density for the LPC versus the lipid tail for Lysolipid_1A_ (Fig. 1e, S2b) and the double conformation seen for the headgroup in the chicken Mfsd2a structure^22^. Second, the flipping of the LPC from Z_A_ to Z_B_ is possible by stabilization of the lipid tail in Chamber_1_ while the headgroup samples multiple binding sites in the open cavity between Z_A_ and Z_B_ (Fig. 2). Because the lipid tail is rigidified in Chamber_1_ and the LPC is translocated from Z_A_ and eventually trapped in Z_B_, the headgroup can reorient from the outward, bent Lysolipid_1A_ to a more linear inward pointing configuration, as observed in Lysolipid_1B_ (Fig. 2, 4c-e). Given these observations, the rotation of the headgroup from the Z_A_ - bound Lysolipid_1A_ to the Z_B_ -bound Lysolipid_1B_ is the next key step in lysolipid flipping. During the process, the headgroup the lysolipid is flipped from the outward to inward-pointing orientation to align the lipid-LPC with the inner membrane leaflet (Fig. 4c-e, S10). Therefore, we propose that the reorientation of the LPC to point the headgroup inward occurs after rotation of the lipid tail outwards and is the next key step in flipping the lysolipid to align with the inner membrane leaflet.

Our ligand- and Na^+^-free drMfsd2a structure likely represents the transporter state in the inward conformation after substrate and cation release (Fig. 4h). In vivo, it is proposed that once the substrate and Na^+^ are released, the transporter resets to an outward opened conformation through a ligand-free occluded state (Fig. 4i-j), completing the transport cycle of Mfsd2a^39^.

### Supplementary Figure legends

**Fig. S1.**
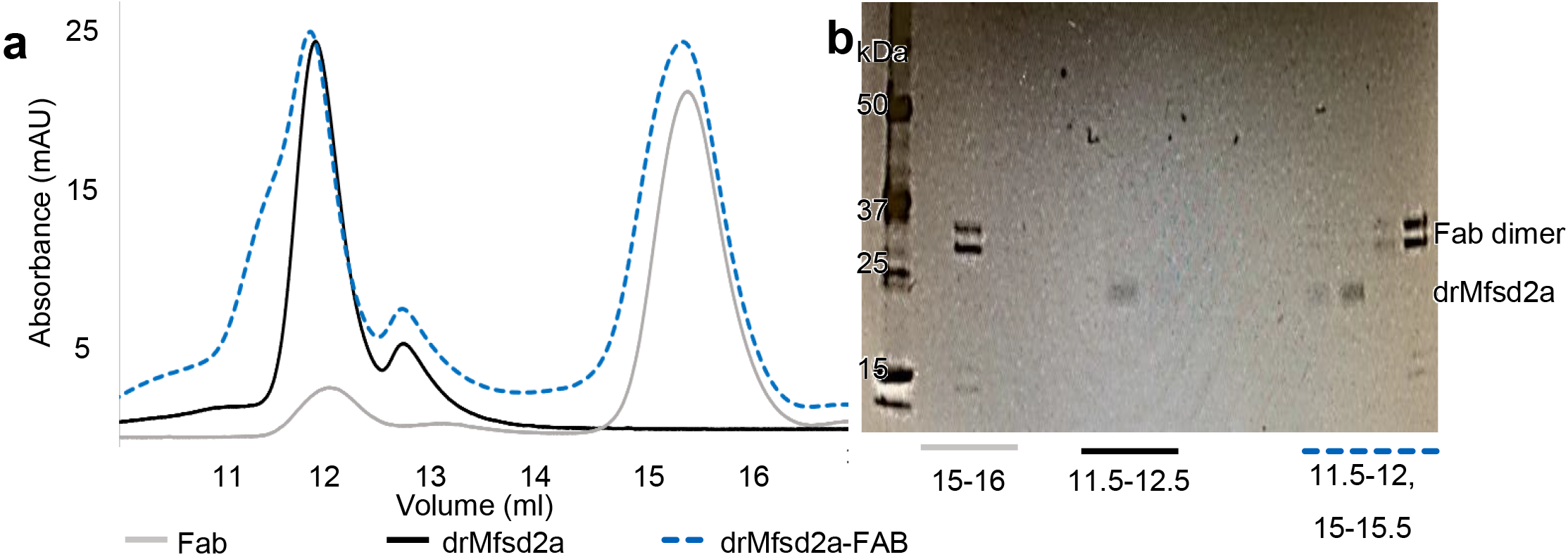
Assembly of the drMfsd2a-FAB complex. **a-b**, Size exclusion chromatography (a) and SDS-PAGE analysis (b) of drMfsd2a -FAB complex formation. Protein was eluted in 0.5 ml fractions. Fractions collected for each sample are indicated.

**Fig. S2.**
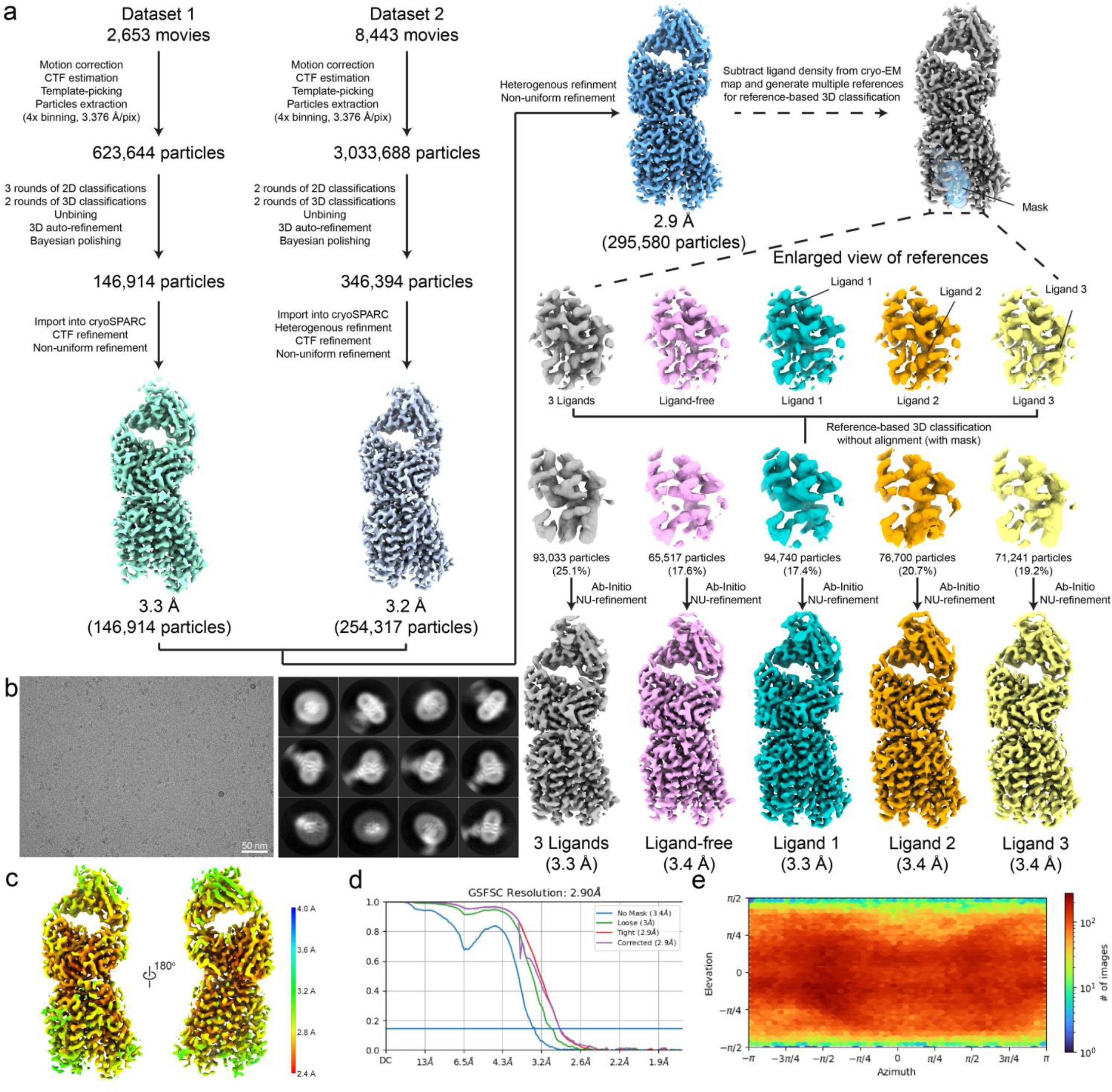
Single-particle cryo-EM data processing for drMfsd2a-FAB complex. **a**, Workflow of cryo-EM image processing of drMfsd2a-FAB. The resolution was reported according to the Fourier shell correlation (FSC) = 0.143 criteria. **b**, A representative cryo-EM micrograph of drMfsd2a-FAB and 2D class averages with a box size of 162 Å. **c**, Local resolution evaluation of the drMfsd2a-FAB map at an average 2.9 Å resolution. **d-e**, Evaluation of the cryo-EM reconstruction of the drMfsd2a-FAB final map with FSC curves (d) and Euler angle distribution plots (e).

**Fig. S3.**
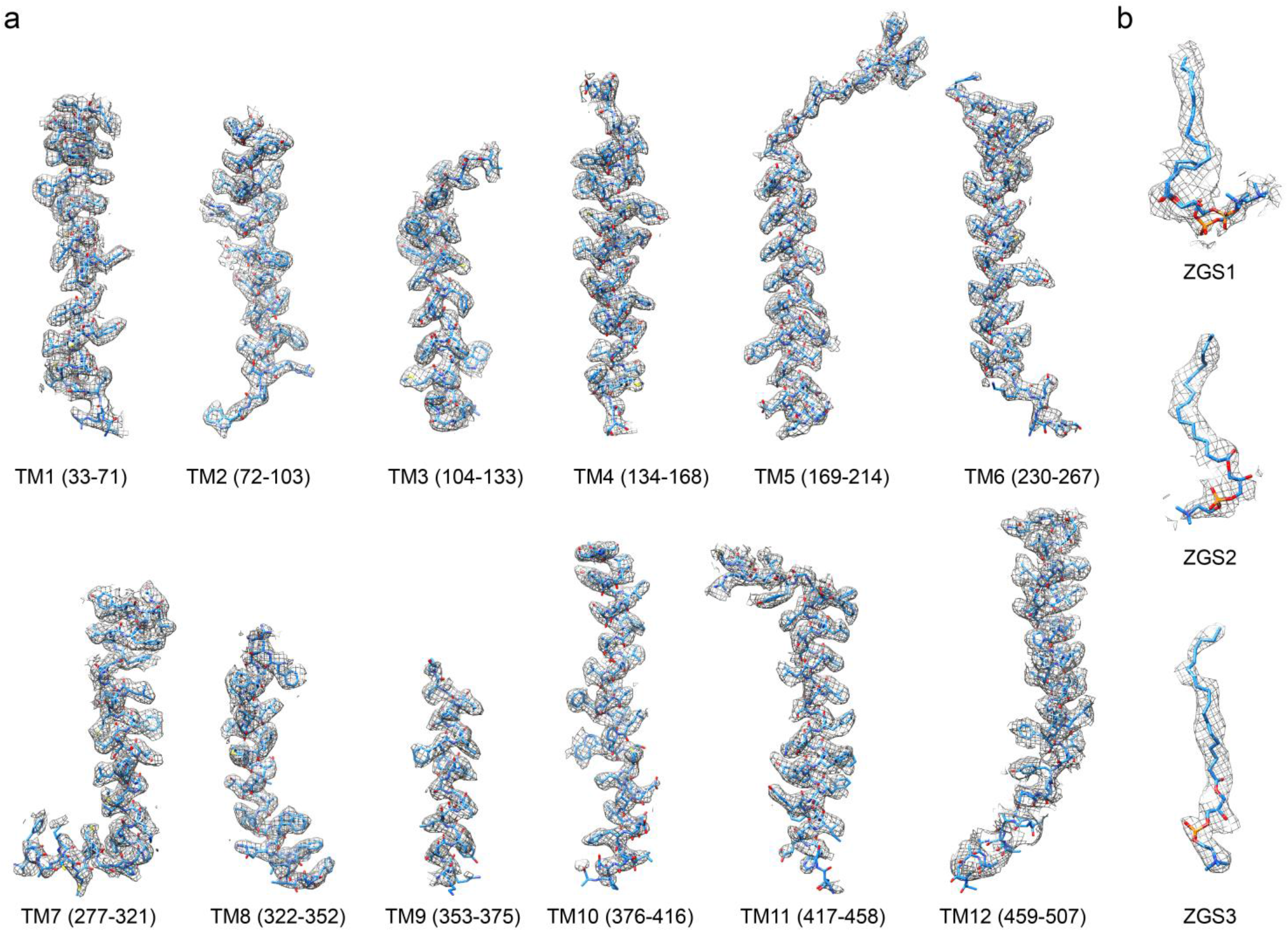
Fit of the drMfsd2a model with cryo-EM density. **a-b**, Cryo-EM densities (mesh) are superimposed on the TMs (a) and ligands (b) of the drMfsd2a model. The model is shown in stick representation.

**Fig. S4.**
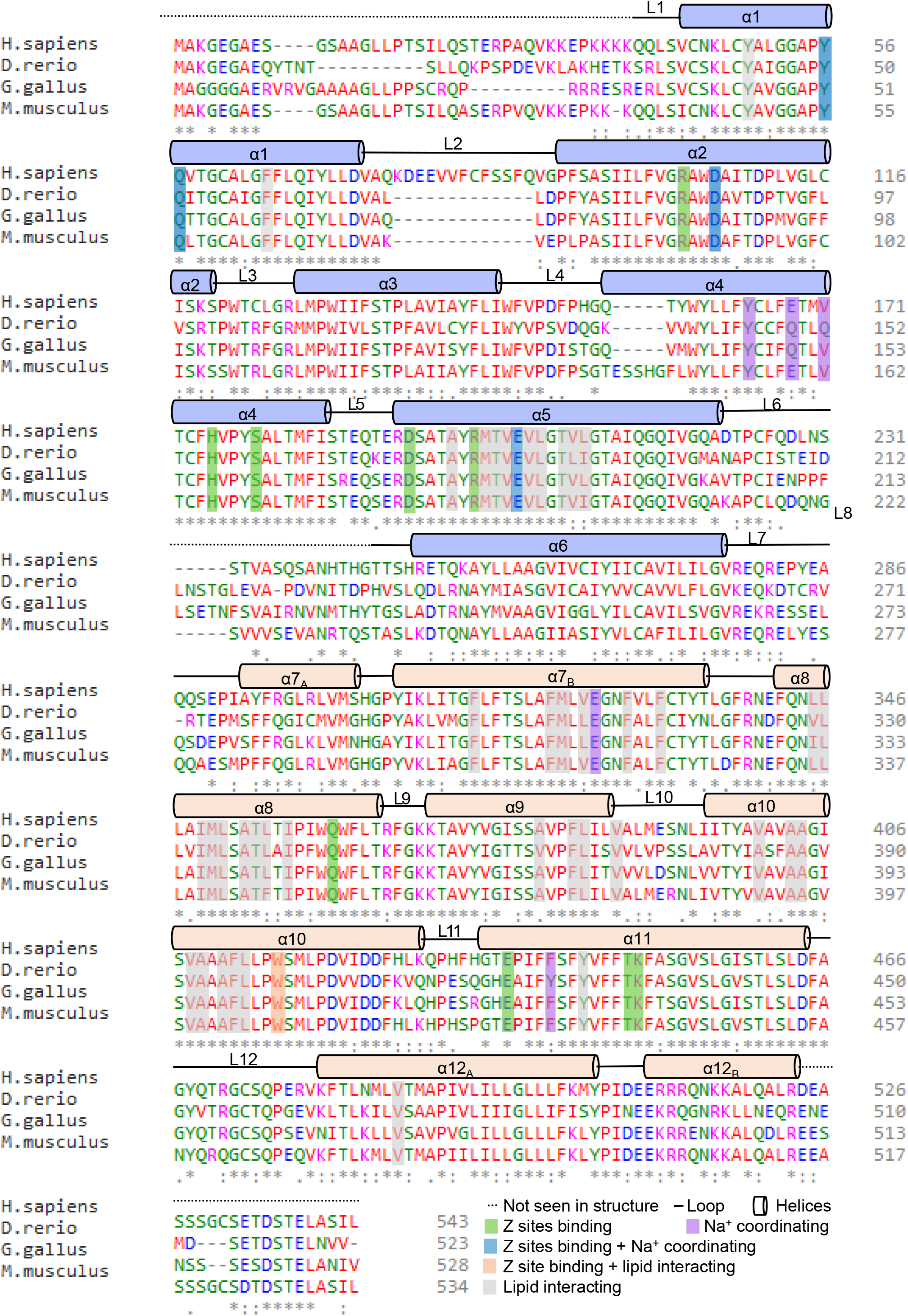
Mfsd2a sequence alignment. Clustal Omega sequence alignment between human (*H. sapiens*, UniProtKB - Q8NA29), zebrafish (*D. rerio*, UniProtKB - Q6DEJ6), chicken (*G. gallus*, (NCBI XM_417826), and mouse (*M. muscuslus*, UniProtKB - Q9DA75) drMfsd2a. Loops and helices are indicated by lines and bars. Residues for Z-site binding, Na^+^ coordinating, and lipid interaction are highlighted in green, purple, blue, wheat and grey, respectively. Unassigned N- and C-terminal domains in dotted line.

**Fig. S5.**
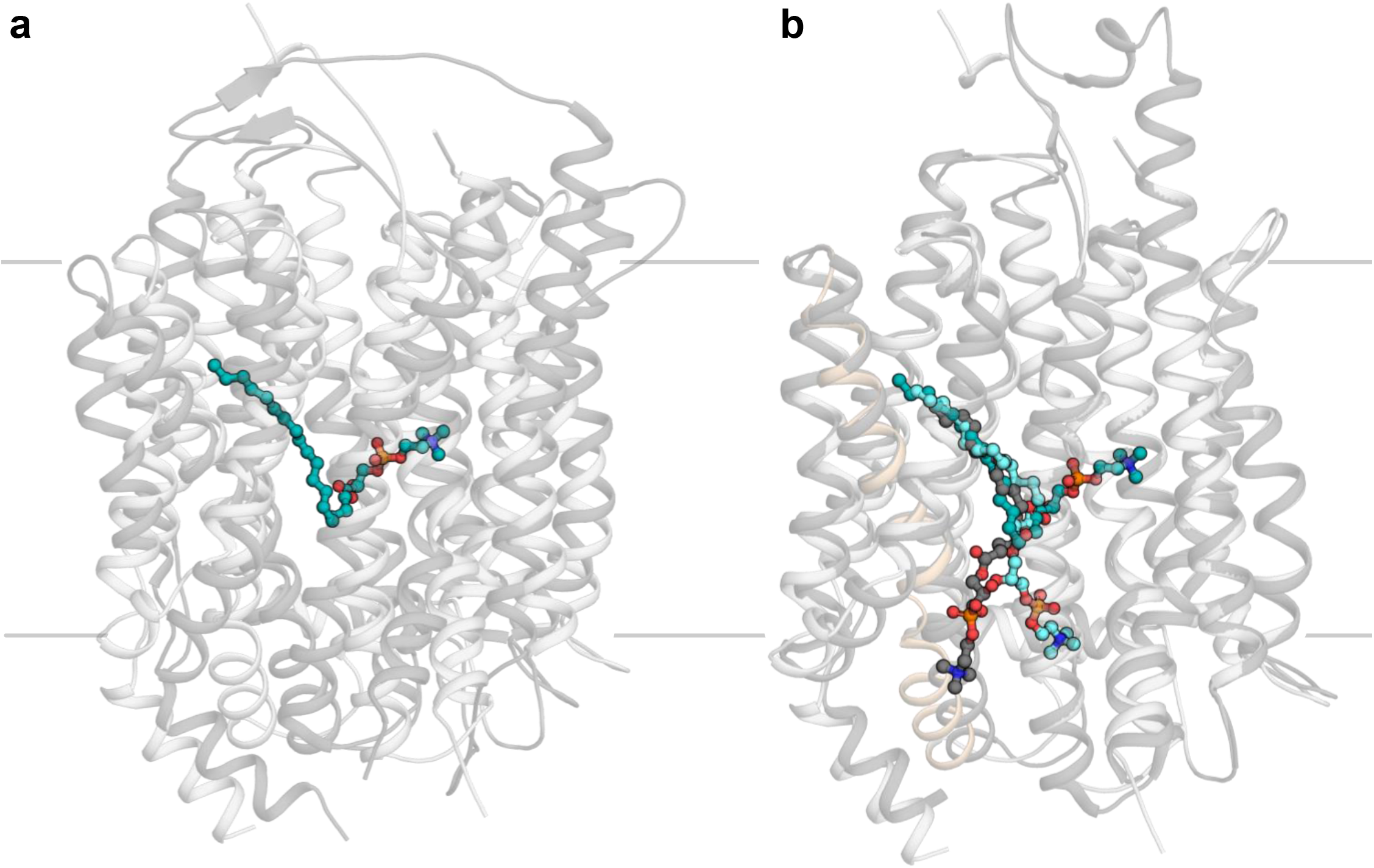
Comparison between the mouse, chicken and zebrafish Mfsd2a structures. **a**, Alignment of mouse^33^ and drMfsd2a. Mouse Mfsd2a^33^ (PDB 7n98) is in dark grey. drMfsd2a is in light grey. Lysolipid_1A_ is in dark cyan. **b**, Alignment of chicken^22^ (PDB 7mjs) and drMfsd2a. Chicken Mfsd2a^22^ is in dark grey. DrMfsd2a is in light grey. Helix 10 of drMfsd2a is highlighted in wheat. ALA-LPC from chicken^33^ Mfsd2a is in grey stick and sphere. Lysolipid_1A-1B_ are in dark and light cyan stick and sphere, respectively.

**Fig. S6.**
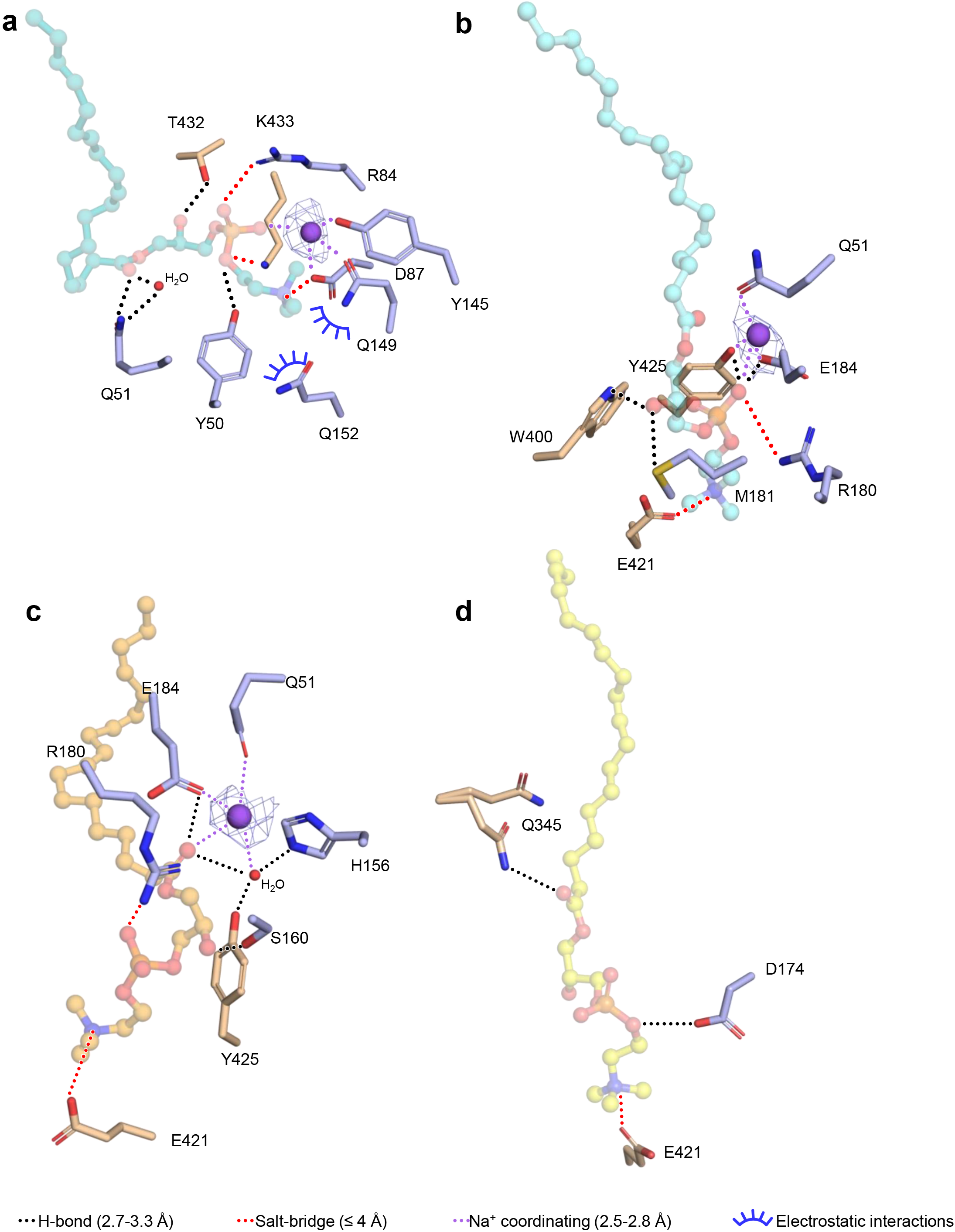
The four LPC binding configurations in drMfsd2a. **a-d**, Alternate views of lysolipid_1A, 1B, 2C, and 3C_ positions. Na^+^ shown in purple spheres. Lysolipids are in stick and sphere. Z-sites and Na^+^ coordinating residues are in stick. Black dotted lines represent H-bonding between 2.6-3.3 Å. Red dotted lines indicate salt bridges with distances ≤4 Å. Purple dotted lines are Na^+^ coordinating functional groups within 2.5-3.2 Å. Blue half circles indicate choline coordinating residues within 3.5 Å. Waters in red small spheres.

**Fig. S7.**
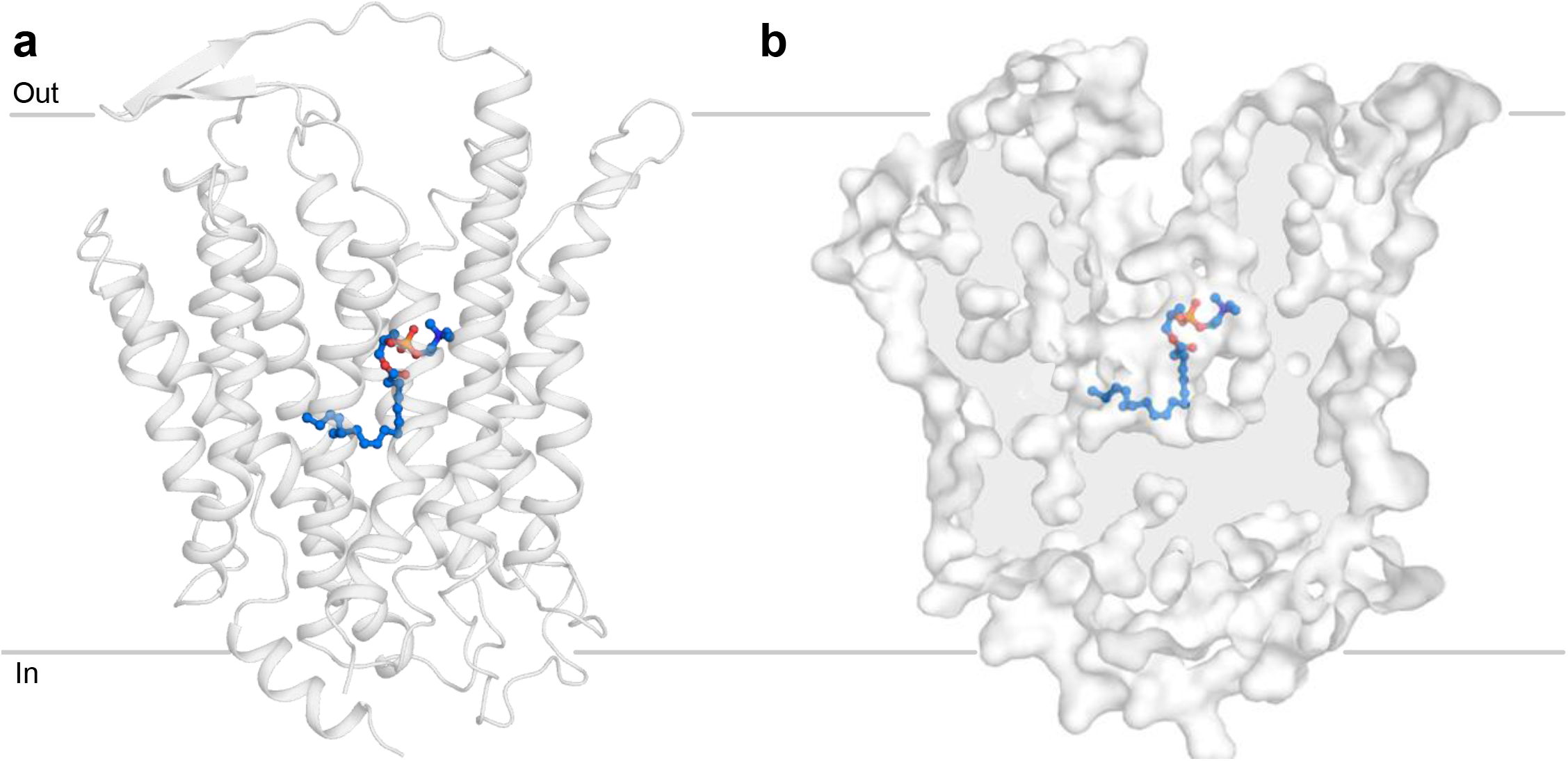
Docking studies of ALA-LPC in the outward-facing mouse Mfsd2a structure. **a-b**, The outward open mouse Mfsd2a structure^33^ (PDB 7n98) with a docked ALA-LPC substrate shown as cartoon (a) and surface (b) representation. Docked ALA-LPC shown as blue stick and sphere.

### Supplementary videos

**Video S1.**
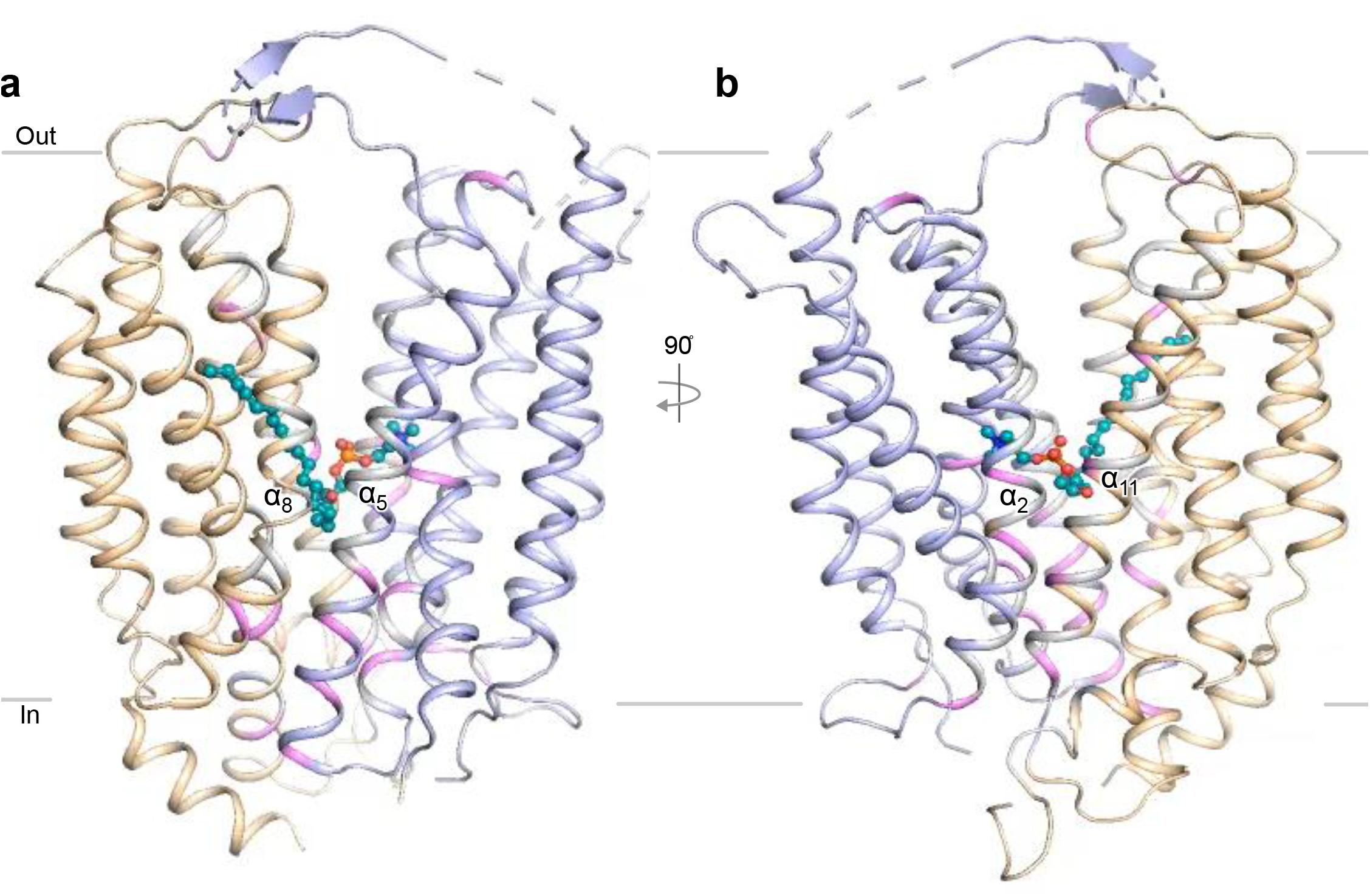
Interdomain contacts and rocker-switch movement between the inward and outward conformation of Mfsd2a. **a-b**, Morphing of the outward mouse^33^ to inward drMfsd2a structure was accomplished by the Morph function in PyMol. Van der Waals interactions (grey) salt bridges (pink), and hydrogen bonds (pink) that make up the interdomain contacts are highlighted. N- and C-domains are in light blue and wheat. The Lysolipid_1A_ is in teal stick and ball.

**Video S2.**
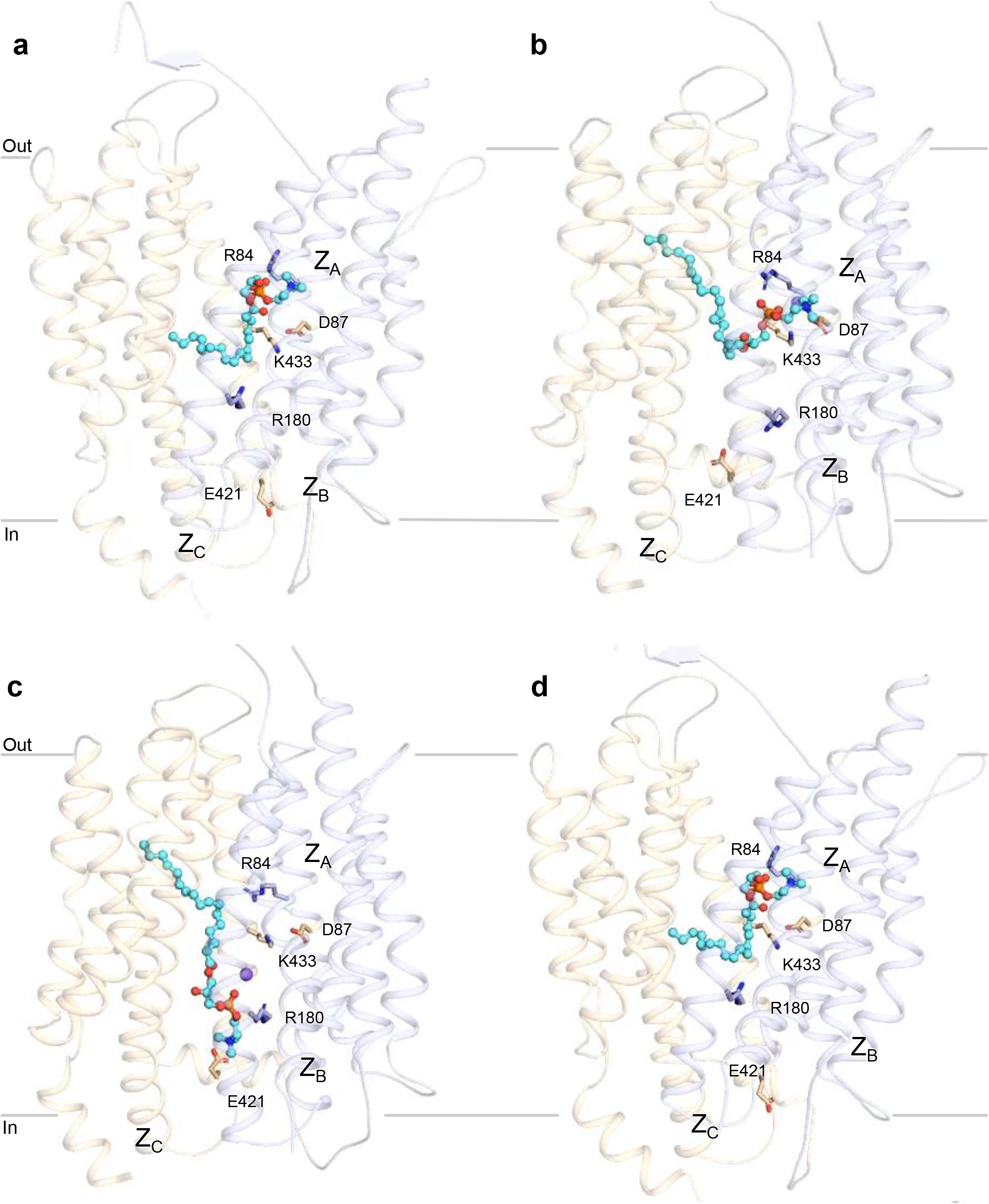
Moving figures of protein conformational changes and lipid flipping, translocation, and release by drMfsd2a. **a**, Conformational changes during the rotation of the lipid tail from the outward-facing mouse^33^ (PDB 7n98) ALA-LPC docked model to the inward-facing drMfsd2a bound with Lysolipid_1A_. **b**, Conformational changes during rotation of the LPC headgroup from the Lysolipid_1A_ to the Lysolipid_1B_ position. **c**, The translocation and cytoplasmic release of the lipid-LPC from the Lysolipid_2B_ to the Lysolipid_3C_ position. **d**, The overall conformational changes from the outward-facing ALA-LPC docked to the release of Lysolipid_3C_ position.

### Supplementary tables

**Table S1.** Cryo-EM data.

|  | Dataset 1 (20210317) | Dataset 2 (20210614) |
| --- | --- | --- |
| Protein concentration | 3 mg/ml | 3 mg/ml |
| Sample volume for EM grid | 3 μl | 3 μl |
| Grid type | QF 400-mesh R 1.2/1.3 Cu | 300-mesh UltrAuFoil 1.2/1.3 Au |
| Plunge freezer | Leica EM GP | Leica EM GP |
| Blotting time (s) | 6 | 6 |
| Blotting temperature ($^{\circ}$C) | 4 | 4 |
| Blotting chamber humidity set (measured) | 95 % (86-91%) | 95 % (86-91%) |
| Microscope | Krios 1 (HHMI Janelia) | Krios 1 (HHMI Janelia) |
| Voltage (kV) | 300 | 300 |
| Camera | K3 (Gatan) | K3 (Gatan) |
| Energy Filter | yes | yes |
| Cs corrector | yes | yes |
| Magnification (x) | 81,000 | 81,000 |
| Physical pixel size ($\text{\AA}$/px) | 0.844 | 0.844 |
| Super-resolution pixel size ($\text{\AA}$/px) | 0.422 | 0.422 |
| Final pixel size used for final maps ($\text{\AA}$/px) | 0.844 | 0.844 |
| Electron exposure ($e^{-}/\text{\AA}^2$) | 50 | 50 |
| Dose rate ($e^{-}/\text{px/s}$) | 9.5 | 9.5 |
| Defocus (μm) | -0.5 to -1.8 | -0.5 to -1.8 |
| Number of total micrographs | 2,679 | 8,443 |
| Number of selected micrographs | 2,653 | 8,443 |
| Number of particles picked | 623,644 | 3,033,688 |
| Number of FAB bound particles | 146,914 | 346,394 |

**Table S2.**
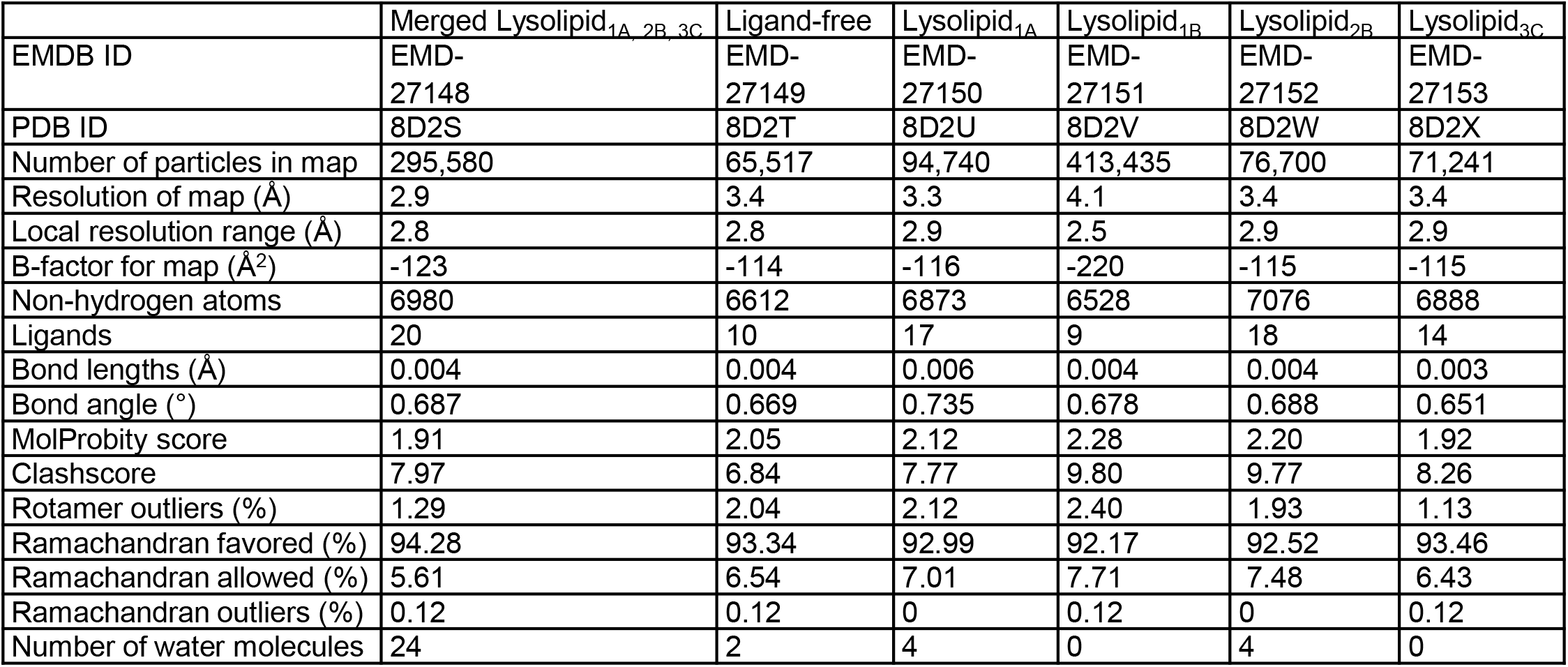
Cryo-EM map and model analysis.

|  | Merged Lysolipid<sub>1A, 2B, 3C</sub> | Ligand-free | Lysolipid<sub>1A</sub> | Lysolipid<sub>1B</sub> | Lysolipid<sub>2B</sub> | Lysolipid<sub>3C</sub> |
| --- | --- | --- | --- | --- | --- | --- |
| EMDB ID | EMD-27148 | EMD-27149 | EMD-27150 | EMD-27151 | EMD-27152 | EMD-27153 |
| PDB ID | 8D2S | 8D2T | 8D2U | 8D2V | 8D2W | 8D2X |
| Number of particles in map | 295,580 | 65,517 | 94,740 | 413,435 | 76,700 | 71,241 |
| Resolution of map (Å) | 2.9 | 3.4 | 3.3 | 4.1 | 3.4 | 3.4 |
| Local resolution range (Å) | 2.8 | 2.8 | 2.9 | 2.5 | 2.9 | 2.9 |
| B-factor for map (Å<sup>2</sup>) | -123 | -114 | -116 | -220 | -115 | -115 |
| Non-hydrogen atoms | 6980 | 6612 | 6873 | 6528 | 7076 | 6888 |
| Ligands | 20 | 10 | 17 | 9 | 18 | 14 |
| Bond lengths (Å) | 0.004 | 0.004 | 0.006 | 0.004 | 0.004 | 0.003 |
| Bond angle (°) | 0.687 | 0.669 | 0.735 | 0.678 | 0.688 | 0.651 |
| MolProbity score | 1.91 | 2.05 | 2.12 | 2.28 | 2.20 | 1.92 |
| Clashscore | 7.97 | 6.84 | 7.77 | 9.80 | 9.77 | 8.26 |
| Rotamer outliers (%) | 1.29 | 2.04 | 2.12 | 2.40 | 1.93 | 1.13 |
| Ramachandran favored (%) | 94.28 | 93.34 | 92.99 | 92.17 | 92.52 | 93.46 |
| Ramachandran allowed (%) | 5.61 | 6.54 | 7.01 | 7.71 | 7.48 | 6.43 |
| Ramachandran outliers (%) | 0.12 | 0.12 | 0 | 0.12 | 0 | 0.12 |
| Number of water molecules | 24 | 2 | 4 | 0 | 4 | 0 |

### Supplementary methods

#### Molecular docking for ALA-LPC in the outward-open conformation of mouse Mfsd2a

<u>The</u> molecular docking of ALA-LPC in the outward-open conformation of mouse Mfsd2a^33^ (PDB 7N98) was calculated with Chimera Autodock Vina tool^51^. The docking was performed within the defined bounding box in the extracellular access of the transmembrane domain, which included a volume of ∼30,000 A^3^ as the search space. Ten docking poses were generated with exhaustiveness of search set at 8, maximum energy difference (kcal/mol) set at 2. The resulting poses yielded scores ranging from 6.2 to -5.7 kcal/mol. The best one was chosen following biochemical rationales.

